# Embodied Navigation: whole-body movement drives path integration in large-scale free-walking virtual reality

**DOI:** 10.1101/2025.11.05.686699

**Authors:** Jonas Scherer, Anabel Kroehnert, Martin Egelhaaf, Norbert Boeddeker

## Abstract

Human navigation relies on combining body cues (vestibular and proprioceptive signals) with visual cues such as optic flow. The weighting and integration of these signals during the continuous tracking of walked distances and angles, known as path integration, remain poorly understood. Previous path integration studies have been limited to small spaces (< 150 *m*^2^) and the influence of complex environments on cue weighting of body and vision cues is still unclear. Here, we conducted the largest-environment free-walking virtual reality navigation study to date in a 45 × 25 *m* facility (1, 215 *m*^2^) using triangle completion tasks in naturalistic environments. We systematically manipulated sensory input across three conditions: natural active walking (full sensory integration), active joystick control (visual cues only), and blindfolded active walking (body cues only). Participants navigated through both sparse fallow land without trees and more complex forest environments with 400 trees. We embedded performance data in a Bayesian cue combination model to analyse the underlying combination mechanism. Our results provide evidence that most participants substantially favour body cues over visual cues in a non-Bayesian combination process, with considerable inter-individual variance in cue dominance strength and side biases. While transitioning from fallow land to forests reduced directional variance, weighting of body and visual information remained constant. These findings advance our understanding of human spatial navigation by demonstrating that body-based cues dominate path integration even in visually rich, large-scale environments, challenging assumptions about optimal Bayesian cue integration in human navigation.

## 1 Introduction

Much like an ancient ship navigator who determines the position using a map, a compass, or the horizon, the human brain continuously processes a complex array of sensory signals to orient and navigate diverse environments from unfamiliar cities, to familiar rooms even in the dark. In the yet largest-environment free-walking virtual reality (VR) navigation study in a 45 *m ×* 25 *m* gym we here investigate how humans weight and combine information from their body and from vision in self-localisation.

Navigation uses two types of cues: external (allothetic) cues from the surroundings and internal (idiothetic) cues from the body[1, 2]. One particularly salient allothetic cue are landmarks that provide reference points through stable visual, auditory, or olfactory markers, and can be used for route planning, orientation, and spatial memory consolidation[3, 4]. While moving, we continuously keep track of idiothetic cues such as body motion, acceleration, and positioning of joints to estimate travelled distances and angles for self-localisation in our immediate environment–a mechanism referred to as path integration[1, 5]. Path integration is based on three sources of sensory information: the vestibular system, proprioception, and optic flow[6]. The vestibular system signals rotational movement through the semicircular canals in the inner ear, which is integrated into head-direction processing[7, 8, 9]. Proprioception feeds into processing of translational movement by providing information about body part locations, muscles, joints, and motor efference copies[9, 10, 11]. Optic flow is the pattern of visual motion perceived during self-movement, where objects in the visual field appear to move in a direction opposite to the observer’s motion[6, 12]. We continuously combine these body (vestibular and proprioceptive) and visual (optic flow) idiothetic cues with allothetic cues like landmarks which allows us to navigate even complex spatial environments[6, 13, 14].

When combining path integration and landmarks it has been shown that internal weighting of landmarks is strongly influenced by various factors including the landmarks’ location[15, 16], distance[4, 17], salience[16], number[16], and spatial stability[18, 19]. Likewise, path integration performance is influenced by the mode of locomotion, i.e. whether participants actively walk or move with a joystick[6, 13], by visual textures in the environment[6], path length[20], and path shape[20, 21], but not by the number of path segments[21]. Cues from landmarks and path integration are combined in a process that considers each cue’s reliability[4, 14, 18, 22]. This process is often explained by Bayesian models of perception[23, 24] assuming the brain to optimally combine noisy sensory information with prior expectations from past experiences[25]. This cue integration of multiple spatial cues within the Bayesian framework of optimality[26, 27] reduces response variability upon the variance achieved by any single cue[16, 18, 22, 28]. While previous research has established how landmarks and path integration are combined[4, 14, 18], the weighting and combination process of visual and body cues within the idiothetic path integration system remains unclear. Moreover, previous path integration studies were limited to small spaces of less than 150 *m*^2^ [4, 6, 14, 18].

In this study, we aim to understand path integration better in a naturalistic, active-walking triangle completion task in the yet largest scale VR study, 45 *m ×* 25 *m* = 1, 215 *m*^2^, by answering the following questions:

1. How are idiothetic body (vestibular and proprioceptive) and visual (optic flow) cues combined and weighted in the path integration process?
2. How is this combination affected by complex environments?
3. What are the implications of an individual’s weighting of body and visual cues in path integration for their navigation performance?

To answer these questions, we use a navigation experiment, the triangle completion task, that engages participants by actively walking through naturalistic fallow land or forests in an indoor sports facility, using a state of the art VR headset (Meta Quest 3). In a triangle completion task, participants are guided along two straight legs of a triangle and asked to freely return to the start location (homing) to complete the triangle (e.g.[6, 10, 16, 20, 21, 29, 30, 31, 32, 33, 34, 35, 36, 37]). To disentangle the contributions of body and vision cues, we compare three experimental conditions: natural active walking (providing full sensory information), active joystick control (visual cues only), and blindfolded walking (body cues only). We embed performance under these conditions in a Cartesian and in a polar Bayesian cue combination model and compare predicted and observed weighting of visual and body cues to analyse the underlying combination mechanism within the path integration system. Further, we alter the number of trees in the environment to investigate the influence of environmental complexity on cue weighting. Since the VR study is essentially a video game, we acquire participants’ gaming experience to analyse its effects on performance and cue weighting.

Most current navigation research is conducted in VR because it enables controlled, yet interactive environments and allows for precise tracking of spatial behaviour[4, 6, 18] [38, 39, 40, 41, 42, 43]. Some studies allow participants to freely walk but due to room-scale limitations only in a comparably small space (e.g. Kearns et al., 2002 [6]: 144 *m*^2^ (VR), Nardini et al., 2008 [14]: 15 *m*^2^ (not VR), Zhao and Warren, 2015 [4]: 25 *m*^2^ (VR), Chen et al., 2017 [18]: 63 *m*^2^ (VR)). Others favour large-scale desktop VR, which has the disadvantage of not allowing active walking and thus omitting vestibular and proprioceptive sensory information[16, 21, 44]. And yet others use large-scale VR on omnidirectional treadmills[20] that have recently been shown to cause at least some unnatural systematic errors in path integration[45]. These methodological limitations leave open the question of whether findings on path integration generalise to navigation in unrestricted, naturalistically large environments.

Although humans can perform path integration using only visual cues, such as random floor and wall textures, body-based cues tend to dominate behaviour in triangle completion tasks when they are available [6]. In line with this, Chance et al. [13] reported poorer performance in a maze navigation task in which participants had to point back to objects they encountered on their walk when participants used a joystick compared to active walking. Ehinger et al. [9] compared triangle completion performance in a virtual starfield and found no behavioural differences between conditions with only vestibular, only proprioceptive, or both cues. Chen et al. [18] found weighting of visual cues to be stronger in an environment with three distinct landmarks than with just one post. However, their study focused on the combination of landmark guidance and path integration. Hence, the influence of different environments on the combination of body and vision cues within path integration, especially with more than three ambiguous landmarks, remains unknown.

Another important influence on path integration is individual differences[46]. Abilities to form mental representations of space vary strongly between individuals[47, 48, 49, 50], due to factors including strategies[51], reference frames[52, 53, 54], age, gender, upbringing environment, and geographic location[55]. In a study including a re-analysis of data from several influential human navigation studies, and our own VR experiment, we previously showed that individuals possess time-persistent biases in path integration [46]. In triangle completion tasks, these manifest as symmetric and asymmetric performance differences between triangles with only left or only right turns. The symmetric error component indicates the bias of over- or undershooting the correct target angle on both sides, and the asymmetric error component reflects a directional bias to one side. Their physiological or neural origin yet remains unclear. Because the experiment was conducted in desktop VR with participants controlling their avatar with the arrow keys, we proposed individual side biases to originate from visual processing rather than from the vestibular system, or motor control. Hence, actively walking individuals who focus on vestibular and proprioceptive cues rather than visual cues, should show smaller side biases. However, Jetzschke et al. [56] demonstrated that directional biases also persist in blindfolded participants performing angle reproduction tasks, suggesting that such biases may not be purely visual in origin. This raises the question of whether side biases in path integration are shaped by the relative weighting of body and vision cues, and to what extent this weighting varies across individuals. A sufficient number of repetitions for left and right triangles in our study allows us to check for individual symmetric and asymmetric biases.

It remains unclear from previous studies whether the above findings on the combination of idiothetic body and visual cues in path integration generalise to naturalistic environments with scales exceeding 150 *m*^2^, whether the presence of allothetic landmarks affects cue weighting, how variable cue weighting is between individuals, and whether cue weighting follows Bayesian integration principles. Here, we conduct a large-scale active-walking VR navigation task to investigate by what mechanism body and vision cues are combined within the path integration process and how the absence or presence of trees influences weighting of cues. We hypothesise that: (1) dominance of body cues will generalise to our large-scale, ecologically enriched VR task; (2) cue combination will follow Bayesian principles, as previously observed for landmarks and path integration[4, 14, 18]; (3) visual cues will be weighted more strongly when trees are present, consistent with earlier findings comparing ‘rich’ and ‘poor’ environments[18]; (4) substantial inter-individual differences will emerge in cue weighting, as we have shown previously for path integration performance [46]; and (5) participants who favour body cues will show smaller side biases, in line with our earlier suggestion that such biases originate from visual processing [46].

## 2 Results

To understand how idiothetic body (vestibular and proprioceptive) and vision (optic flow) cues are combined within the path integration process and how the presence of a cluttered environment affects their combination, we let *N* = 13 participants complete active free-walking VR triangle completion tasks in a 1, 215 *m*^2^ gym. To assess path integration, participants walked along two legs of a triangle and then freely returned to the start location (see methods ‘Virtual reality triangle completion task’). Besides cue combination trials with both body and vision cues, single cues were isolated in a body condition, in which participants were blindfolded and guided only by controller vibrations, and in a vision condition, in which participants controlled their avatar by joysticks without proprioceptive, vestibular, or efferent motor input (see methods ‘Virtual reality triangle completion task’ for details). This set of conditions allowed us to calculate relative cue weighting of body and vision cues in the path integration process (see methods ‘Bayesian cue combination models’). It was completed by all participants in a fallow land environment without trees, isolating the path integration system as guidance mechanism, and additionally in forests with 400 ambiguous trees to assess the influence of more complex environments on cue weighting within the path integration system. Observed task performance is quantified by position error (Euclidean distance between response and target), distance error (positive: walking too far; negative: not walking far enough), and direction error (positive: turning too far; negative: not turning far enough)(see methods ‘Error calculations’), which are later compared to predictions from Bayesian cue integration models in either a 2D Cartesian coordinate system (for position error) or a polar coordinate system (for distance and direction errors).

### 2.1 Performance errors are larger for vision than for body condition

Navigation performance is strongly influenced by the availability of body and vision cues, as reflected in participants’ homing trajectories under conditions with differing sensory information (Fig. 1A and B). There is large variance in responses between participants in the vision, body, and combination condition. In the combination condition, the response endpoints span the entire gym. In contrast, in the body condition, participants tend to walk shorter distances, with endpoints covering about half of the gym. In the vision condition, response endpoints go beyond the dimensions of the physical gym.

**Figure 1:**
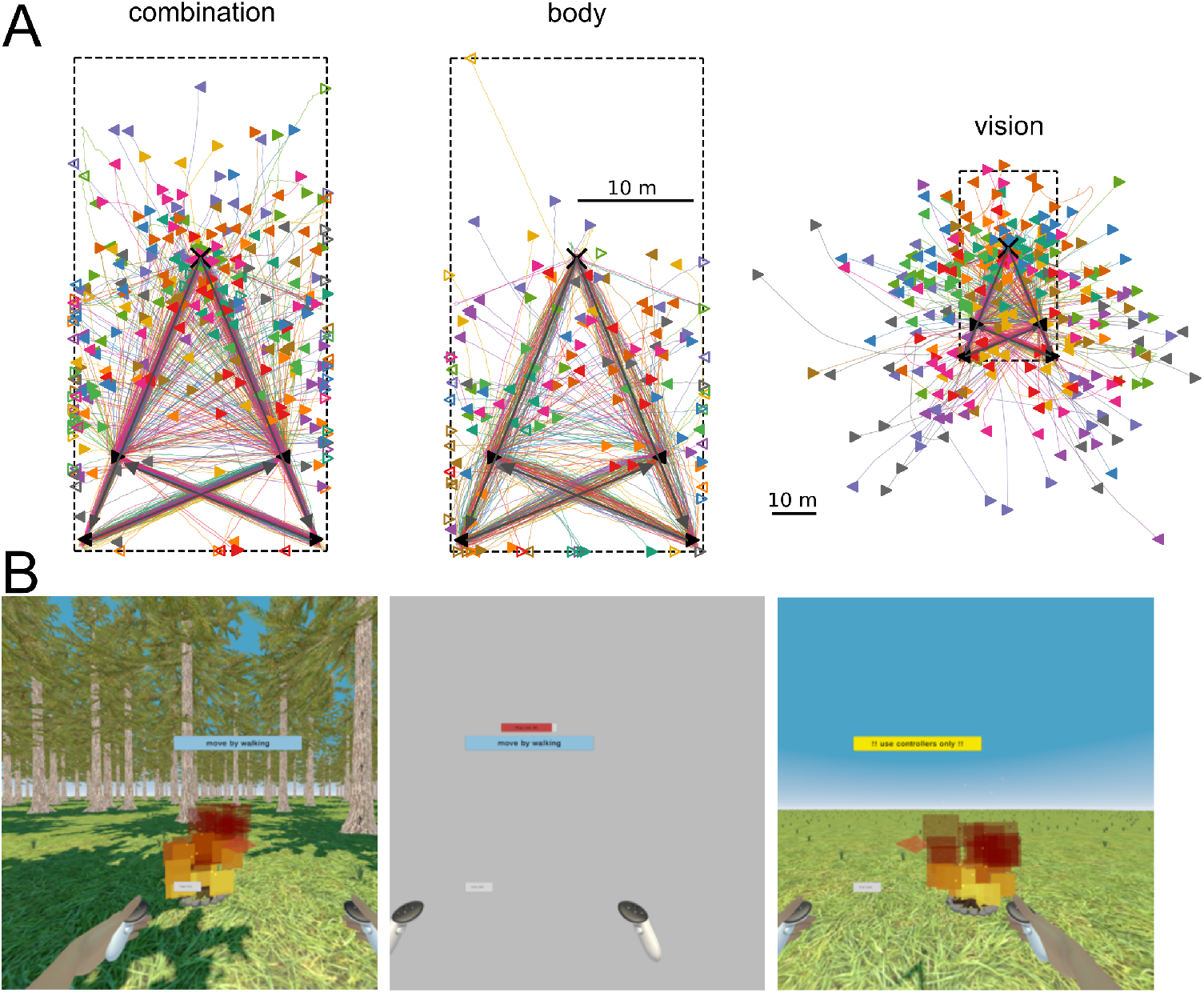
(A) Full homing trajectories for all participants in combination, body, and vision condition with screenshots taken in the above condition (B) either in the 400 tree (left, left-eye view) or the 0 tree environment (right, right-eye view). (A) In each trial, participants walked along two legs of a triangle (grey arrows) indicated by firewood waypoints (black triangles) and then freely returned to the then invisible target location (black cross) on a homing trajectory (coloured lines and coloured triangle markers). Waypoints are collected along a left-hand triangle path (left pointing triangle markers) or on a right-hand triangle path (right pointing triangle markers). For simplicity only one left-hand and one right-hand triangle is shown, and trials from the other target location and gym half are mirrored to the displayed side (Fig. 9). The gym dimensions are indicated by dashed black lines. Trajectories of trials in which participants got too close to the wall and were stopped are indicated by outlined markers at the gym boundary. In the vision condition participants could walk their avatar outside the physical gym dimensions. Homing trajectories of each participant are displayed in the same colour. Scales are given by 10 *m* scale bars. (B) Screenshots contain target campfire, a blue banner saying ‘move by walking’, a yellow banner saying ‘!!use controllers only!!’, a red guidance arrow, a grey trial progress bar, a red timeout bar, and VR arms with controllers.

While in the vision condition participants are not stopped when navigating outside the gym dimensions, in the body and combination condition they are stopped when getting too close to a wall. Respective trials are considered as ‘out-of-bounds’ and responses are considered only for directional error calculations, assuming that participants walk overall straight on their homing path, but are removed for any distance and 2D position calculations. Out-of-bounds trials make up 28% of trials in the body condition and 23% in the cue combination condition with no significant difference between them (paired permutation difference test: *ci* = [−0.047, 0.014], *p* = 0.262)(Fig. 2).

**Figure 2:**
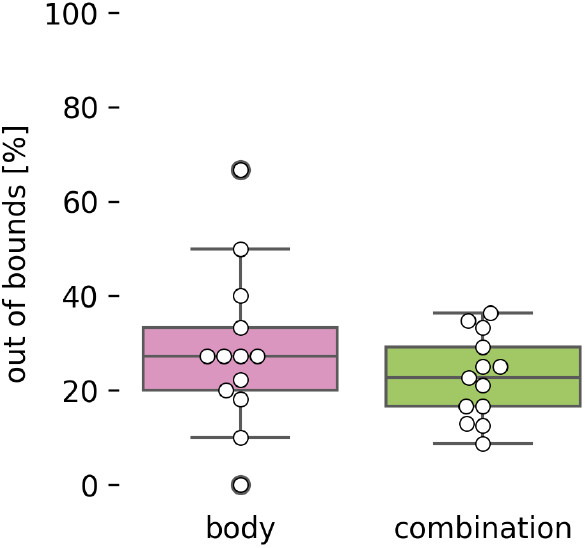
Percentage of trials per condition in which participants were stopped because they walked too close to a wall; these trials were marked ‘out-of-bounds’. Out-of-bounds trials are considered only for directional error calculations, assuming that participants walk overall straight on their homing path, but are removed for any distance and 2D position calculations. Data points represent single participants.

Homing trajectories differ depending on the sensory information available in each condition (Fig. 3). ANOVAs and post-hoc non-parametric paired permutation difference tests reveal that straightness is significantly (*F* = 6.1, *p* = 0.005) lower in the body (0.89) than in the combination condition (0.96)(Cohen’s *d* = 2.1), and that trajectory length (*F* = 14.2, *p <* 0.001) and average speed (*F* = 45.7, *p <* 0.001) are highest in the vision condition 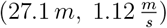 and lowest in the body condition 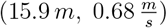(body vs vision: trajectory length: *d* = 1.73, average speed: *d* = 2.42) (Fig. 3). Differences between conditions underline that participants navigate differently depending on available sensory information.

**Figure 3:**
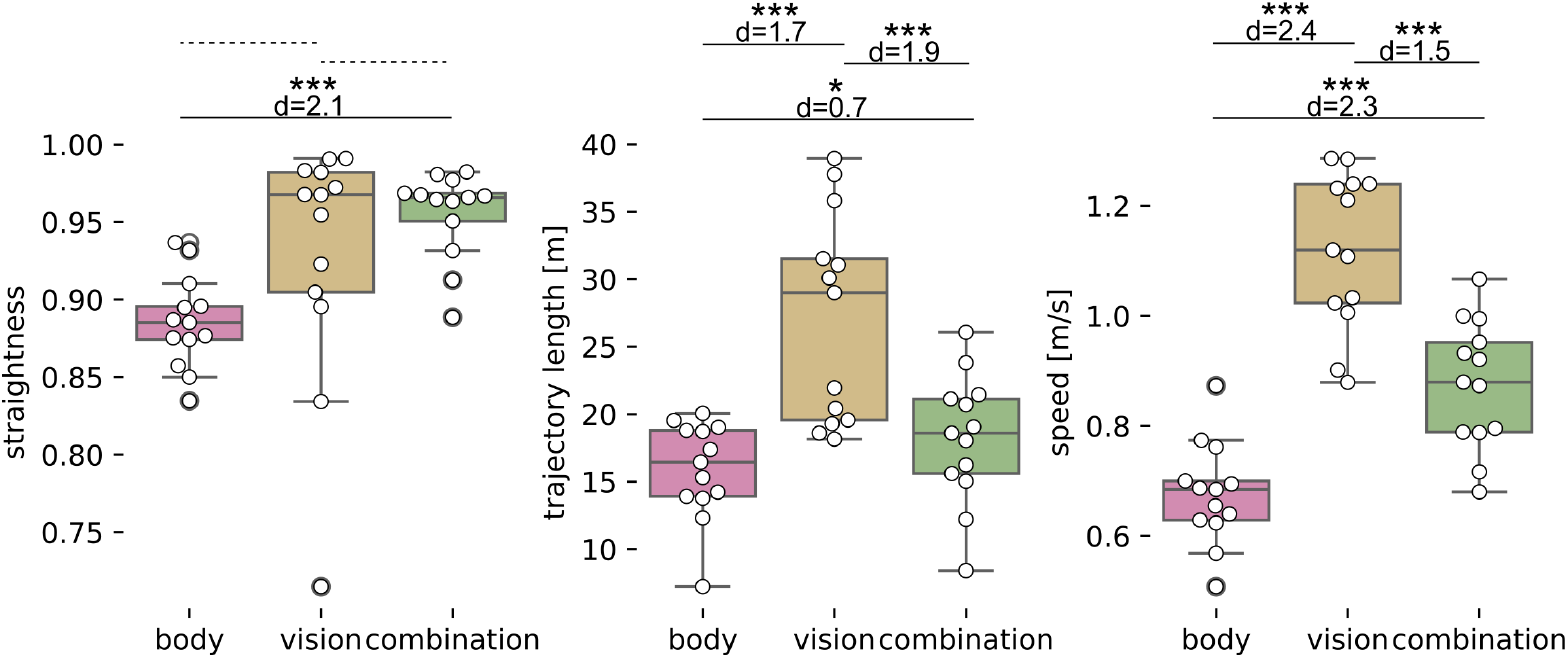
Straightness (walked distance / bee line) (A), trajectory length (B), and average speed (walked distance / total duration) (C) per condition. Trajectories for right and left-hand triangles as well as for trials with 0 or 400 trees are pooled. Significance code (∗ : *p* < 0.05, ∗∗: *p* < 0.01, ∗∗∗: *p* < 0.001) is based on Bonferroni corrected paired permutation difference tests. Effect sizes are indicated by Cohen’s d. Non-significant comparisons are indicated by dashed lines. Data points represent single participants’ means.

Performance varies between the different sensory conditions (body, vision, combination) and environments (0 or 400 trees) in position error, distance error, and direction error (Fig. 4). To assess the effects of condition and number of trees on performance errors we calculate linear mixed models (LMMs) in which position, distance, or direction error is predicted from the fixed categorical effects ‘condition’ and ‘number of trees’, their interaction, and a random ‘participant’ effect (*error* ∼ *condition* ∗ *n*_*trees*_ +(1|*participant*))(for effect sizes see Fig. 4, for details on LMMs see supplement ‘**??**’).

**Figure 4:**
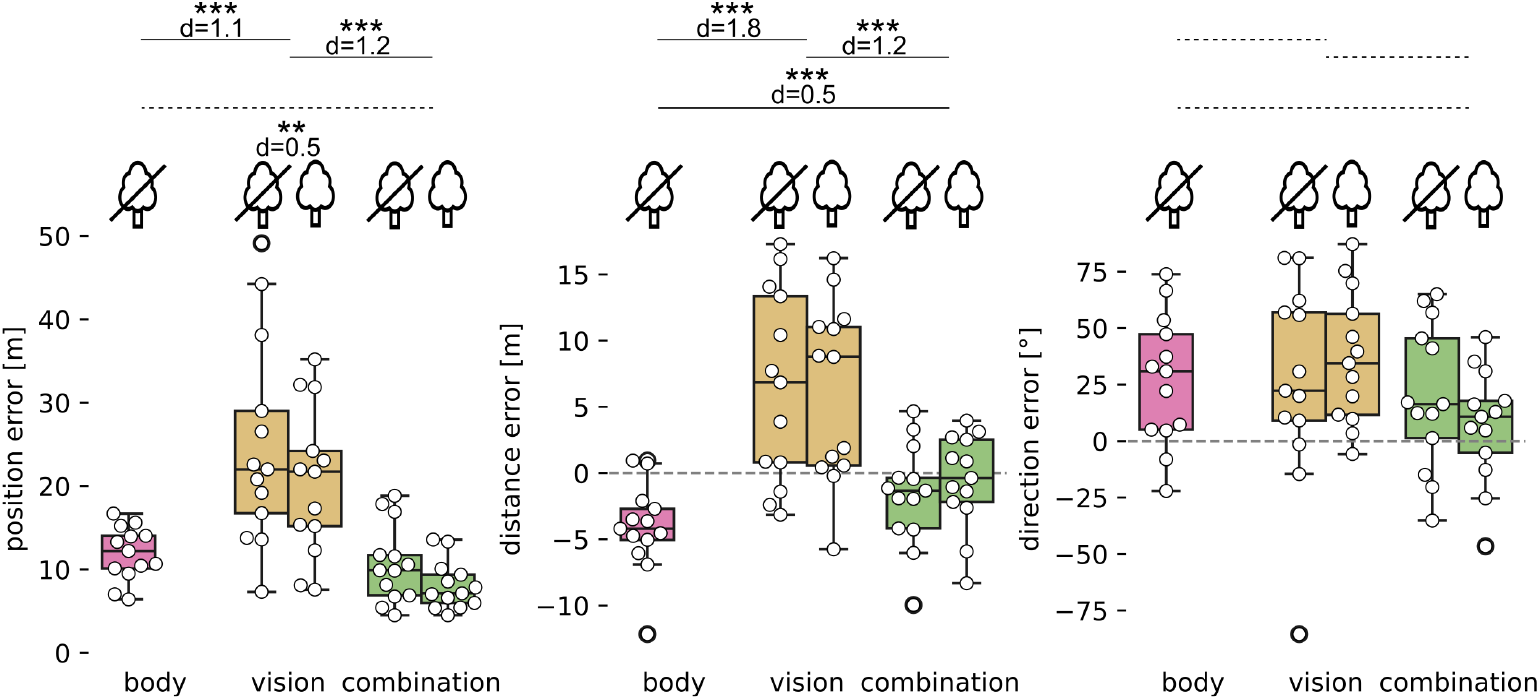
Position (A), distance (B), and direction error (C) in body, vision, and combination condition, and environments (0 trees: crossed out tree icon; 400 trees: uncrossed tree icon). Direction errors correspond to a standardised right-hand triangle. Data points represent single participants’ means. Significance code (∗: *p* < 0.05, ∗∗: *p* < 0.01, ∗∗∗: *p* < 0.001) is based on Bonferroni corrected contrasts of estimated marginal means of LMMs using ‘emmeans’ package (1.11.1) in R (4.5.0). Effect sizes are indicated by Cohen’s *d*. Non-significant comparisons are indicated by dashed lines.

LMMs and post-hoc contrasts of estimated marginal means (see Fig. 4 caption for details) reveal that position errors are significantly higher in the vision condition (22.5 *m*) than in the body condition (12.4 *m, d* = 1.1) or combination condition (10.6 *m, d* = 1.2, see supplement Tab. S1-2), and that distance errors are larger in the vision condition (6.3 *m* overshoot) than in the combination condition (1.3 *m* undershoot, *d* = 1.2) which are in turn larger than in the body condition (4.5 *m* undershoot, *d* = 0.5, see supplement Tab. S3-4). Direction error is neither significantly affected by condition nor by the number of trees (see supplement Tab. S5-6).

Variance in error measures is quantified by the standard deviation (sd) of position error (in *m*), distance error (in *m*), and direction error (circular sd in ° estimated from concentration coefficient *κ* of von Mises distribution; see section ‘Body cues are weighted more than vision cues’). For error variance, equivalent LMMs and post-hoc contrasts of estimated marginal means (see Fig. 5 caption for details; integration and alternation model predictions are presented later in section ‘Body cues are weighted more than vision cues’) reveal that standard deviation (sd) in 2D Cartesian space (see supplement Tab. S7-8), and in distance (see supplement Tab. S9-10) are significantly higher in the vision (Cartesian 2D: 14.1 *m*; distance: 5.9 *m*) than in the body (Cartesian 2D: 5.3 *m, d* = 3.4; distance: 2.3 *m, d* = 2.4) or combination condition (Cartesian 2D: 5.9 *m, d* = 3.1; distance: 3.2 *m, d* = 1.9). Direction sd does not differ significantly between the body (30.1°) and vision (34.7°) condition, and is lowest in the combination condition (24.9°, vision vs combination: d = 1, see supplement Tab. S11-12). Across participants, response variance was lowest in the body condition for both 2D Cartesian and distance estimation. For direction estimation, however, 4 out of 13 participants showed lower variance in the vision condition than in the body condition. According to Bayesian cue integration principles, combining body and vision cues should reduce response variance in the combination condition relative to either single-cue condition (vision or body). Yet, considering each participant’s lowest single-cue variance, the combination condition does not yield a significant reduction in response variance for 2D Cartesian, distance, or direction estimation.

**Figure 5:**
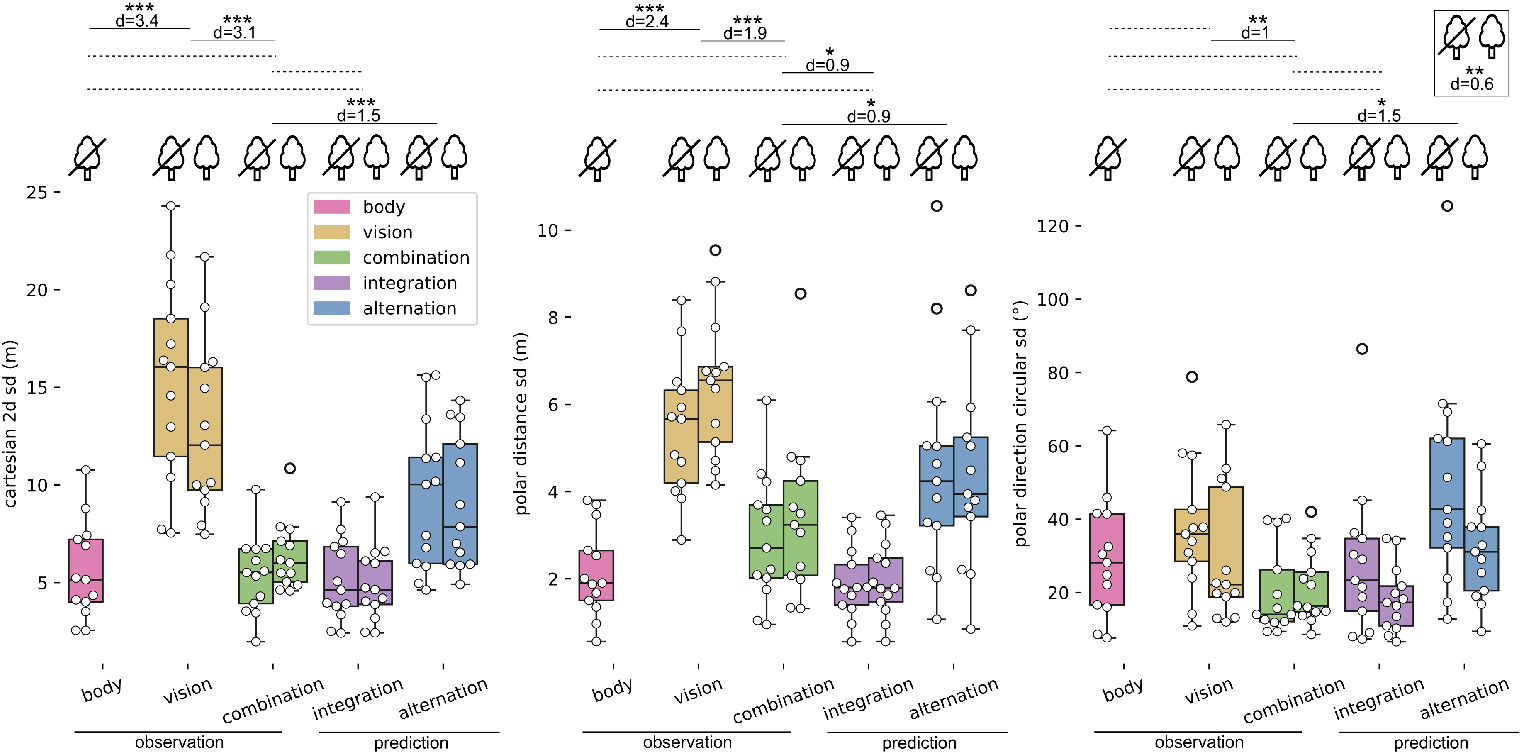
Observed variability in body, vision, and combination condition, and predicted variability in integration, and alternation model for the Cartesian 2D model (A, sd in *m*), the polar distance (B, sd in *m*), and the polar direction model (C, circular sd *κ*^−1^ in) separated for environments (0 trees: crossed out tree icon; 400 trees: uncrossed tree icon). Data points represent single participants’ variance measures. Significance code (∗: *p* < 0.05, ∗∗: *p* < 0.01, ∗∗∗: *p* < 0.001) is based on Bonferroni corrected contrasts of estimated marginal means of LMMs using ‘emmeans’ package (1.11.1) in R (4.5.0). Effect sizes are indicated by Cohen’s *d*. Non-significant comparisons are indicated by dashed lines.

In summary the LMMs and respective effect sizes indicate a large effect (1.1 ≤ *d* ≤ 1.8) of condition on accuracy (mean) in position and distance (Fig. 4A and B), a very large effect (1.0 ≤ *d* ≤ 3.4) of condition on precision (variance) in all error measures (Fig. 5), and a moderate effect (*d* = 0.6) of the number of trees on precision in direction (Fig. 5C). Both the sensory condition and the presence of trees changes how participants navigate in our experiment suggesting that participants show distinct preferences for body or vision cues. In contrast to Bayesian cue integration principles, response variance is not reduced in the combination condition relative to either single-cue condition.

### 2.2 Body cues are weighted more than vision cues

Based on the navigation performance of participants in the single-cue and combination conditions we analyse weighting of body and vision cues within the path integration process. We infer individual preferences for certain cues and assess whether the combination of body and vision cues follows Bayesian cue integration principles or is based on a different mechanism.

We calculate Bayesian cue combination models separately for each participant and for each number of trees (0 or 400). We fit models in a Cartesian coordinate system, following the procedure described by Chen et al. [18], and in a polar coordinate system with separate distance and direction components based on Murray and Morgenstern [57] and Zhao and Warren [4] (see also [16]). For the Cartesian and distance model, mean and sd of a 2D or 1D Gaussian are fitted, respectively. For the direction model, we fit a von Mises distribution (circular normal distribution) with mean direction and concentration coefficient (as a rule of thumb: 1/circular sd)(see methods ‘Bayesian cue combination models’ for details). Based on the Cartesian and polar model we can assess the importance of body and vision cues separately for 2D, distance, and direction estimation and compare observed cue weights with their Bayesian cue combination predictions. While the Cartesian model treats path integration as estimating a single 2D endpoint location, the polar model assumes that the brain separately tracks distance and direction—reflecting a more embodied, movement-centered representation of space. Predicted cue weight, in a Bayesian framework, corresponds to its cue’s relative reliability (1/variance) compared to the total reliability across all cues. Observed cue weights correspond to the relative proximity of the mean combination response to either single-cue response (vision or body). We compare fits of Bayesian integration models (cues are combined by a weighted sum) and Bayesian alternation models (on each trial, a participant relies on either the body or vision cue alone, with the likelihood of choosing each cue determined by its weight) to assess the underlying cue combination process. All models are fitted to responses that are standardised to a right-hand triangle so that the homing start location corresponds to the origin and the target direction aligns with the positive y-axis (Cartesian) or angle zero (polar).

While for some participants the Cartesian and polar Bayesian models fit the observed response data in the combination condition nicely (Fig. 6A) they deviate distinctly for others (Fig. 6B). Most participants do not show distinct differences in their responses between the environment with 0 or 400 trees. Thus, we show representative responses and models only for the 0 tree environment.

**Figure 6:**
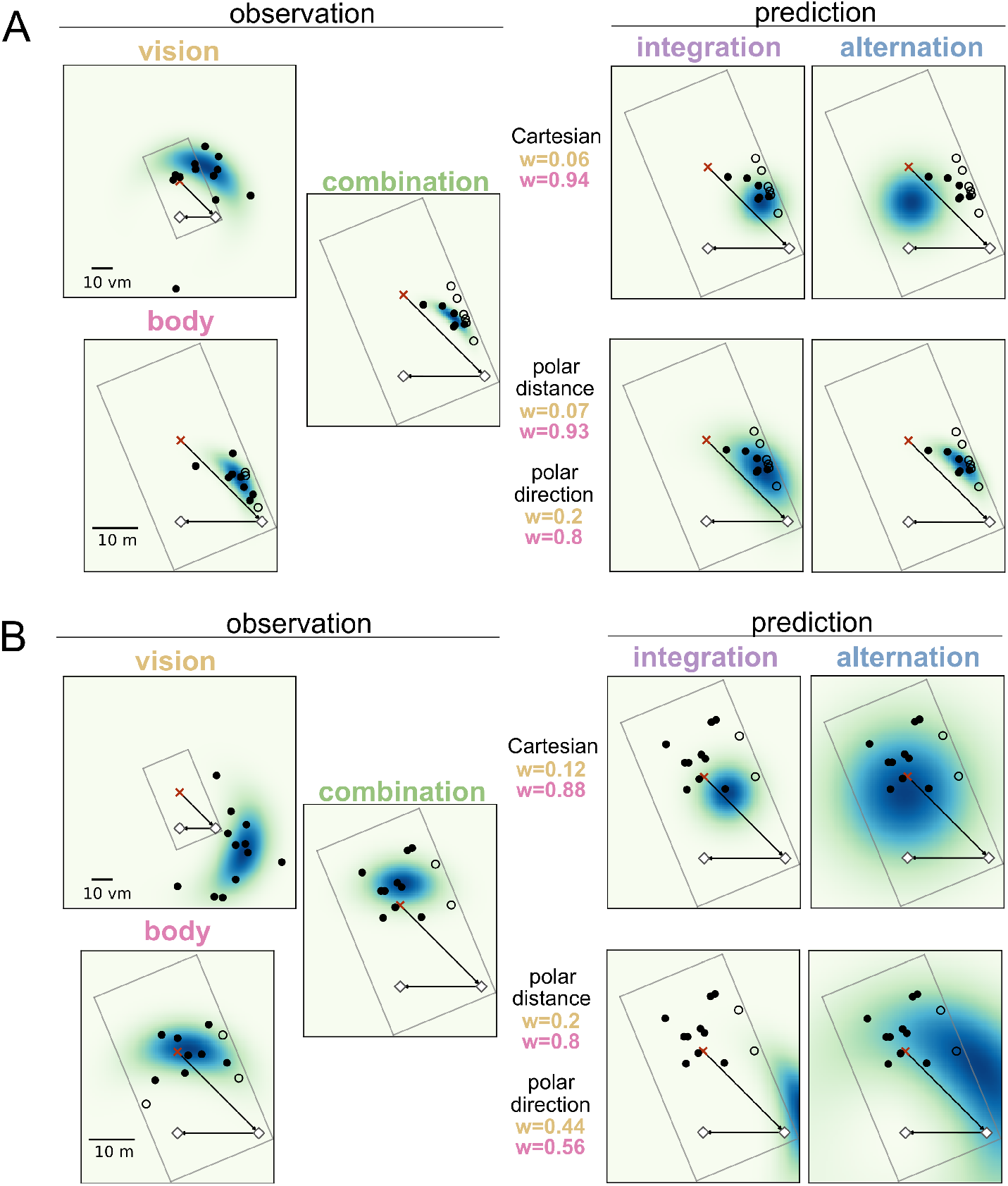
Predictions from Bayesian cue combination models fit observed responses for participant A but not for participant B in a Cartesian and a polar coordinate system. Cue combination models assign weights (*w*) to vision (yellow) and body (magenta) cues based on the cue’s relative reliability. Polar fits are composed of a distance and a separate direction fit with separate cue weighting (see methods ‘Bayesian cue combination models’ for details). Models are fit assuming either cue integration (weighted sum of single cues) or cue alternation (alternation between single cues with weighted probability). Observed responses in the vision, body, and combination condition are overlaid with polar MLE distributions fitted on the observed data. All displayed panels show data from the environment with 0 trees. There are only small performance differences between the environments with 0 or 400 trees.

The above described differences in performance error variance between conditions (Fig. 5)(section ‘Performance errors are larger for vision than for body condition’) can be translated into predicted cue weights in the Bayesian cue combination models. Namely, larger response variance in vision trials than in body trials cause higher predicted weighting of body cues within the path integration system because body cues are more reliable in our navigation task. Accordingly, the above presented main effect of condition on 2D Cartesian, distance, and direction sd, as well as the main effect of number of trees on directional variance imply corresponding effects on cue weighting. Bayesian cue integration predicts that response variance in the combination condition should be lower than in any single-cue condition; as shown above, however, this reduction is not observed in our experiment (Fig. 5). Instead, response variances in the single-cue conditions indicate that body cues, rather than vision cues, predominantly govern responses in the combination condition across participants for 2D position and distance estimation (but not for direction estimation). Furthermore, contrary to Bayesian cue integration principles, the observed weights for visual cues do not correlate with the predictions from the Bayesian models in any domain–2D Cartesian, distance, or direction (Fig. 7B). This indicates that participants did not combine visual and body cues in their path integration system according to Bayesian weighting. Therefore, we consider the observed cue weights more informative than the predicted weights.

**Figure 7:**
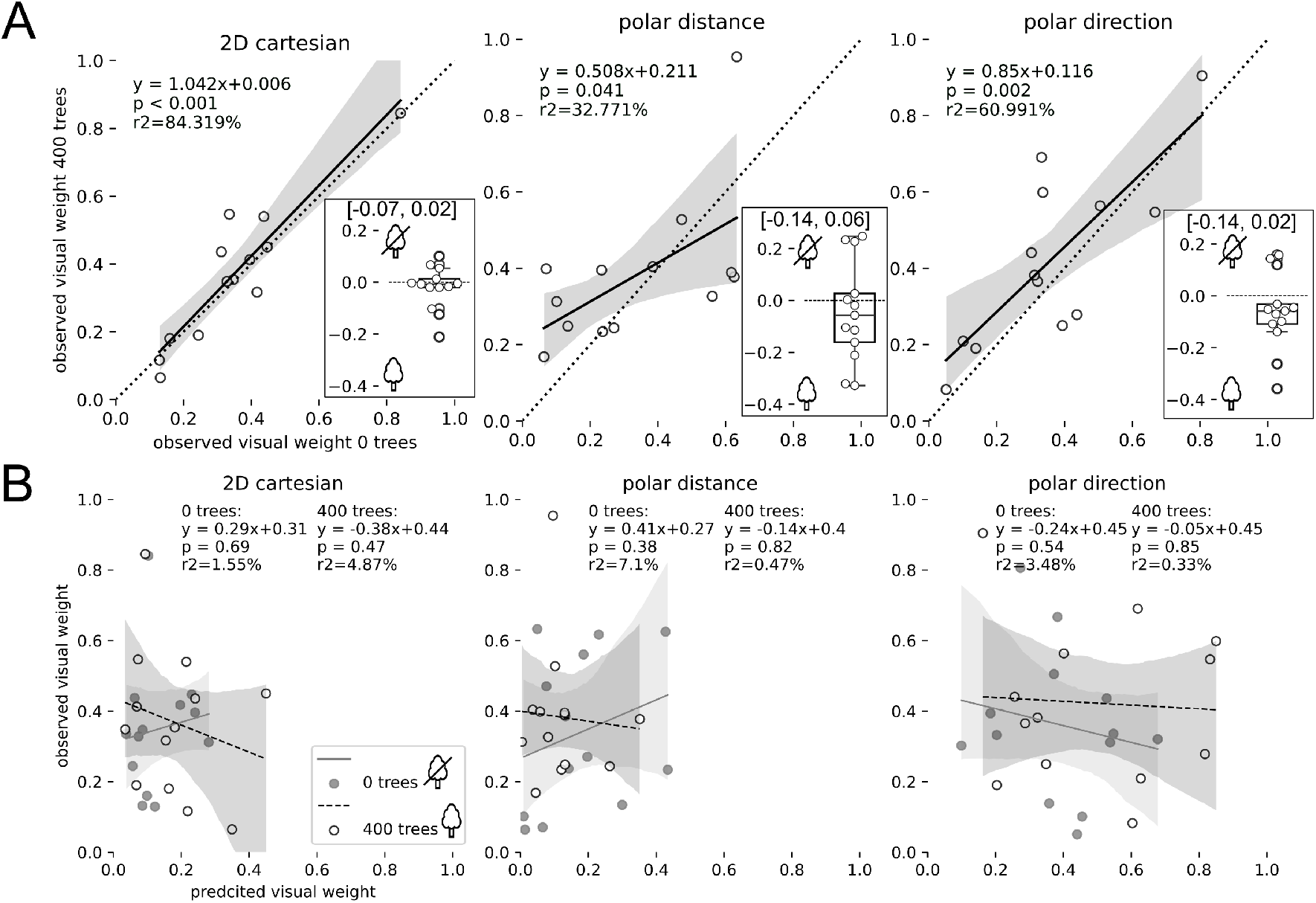
Vision weights correlate between environments with 0 trees and with 400 trees. (A) Differences in vision weights (0 − 400) between the 0 and 400 tree environment are not significantly different from zero (one-sample bootstrap hypothesis testing) and are presented in inserts together with their 95% − *ci*. The same applies to body weights because *w*_*body*_ = 1− *w*_*vision*_. Correlations are described by their 95% − *ci* (shaded area), regression equation (*y*), *p*-value (*p*), and coefficient of determination (*r*^2^). Data points represent single participants. (B) Observed cue weights do not correlate with predictions from Bayesian cue combination models. Models were fit separately for environments with 0 trees (solid line, filled dots) or 400 trees (dashed line, unfilled dots).

Observed cue weights are higher for body than for vision cues across participants in 2D Cartesian (permutation difference test: *mean* = −0.49, *p* < 0.001, 95% − *ci* = [−0.05, 0.13], *d* = 2.4), distance (*mean* = −0.49, *p* < 0.001, 95% − *ci* = [−0.07, 0.07], *d* = −2.2), and direction estimation (*mean* = −0.17, *p* = 0.031, 95%−*ci* = [−0.08, 0.002], *d* = −0.68)(Fig. 7A). Nevertheless, mean weight across participants assigned to vision cues also proves to be significantly different from zero in 2D Cartesian (*mean* = 0.256, bootstrapping hypothesis testing: *p* < 0.001, 95% − *ci* = [0.2, 0.31], *d* = 2.5), distance (*mean* = 0.25, *p* < 0.001, 95% − *ci* = [0.19, 0.32], *d* = 2.2), and direction estimation (*mean* = 0.42, *p* < 0.001, 95% − *ci* = [0.36, 0.48], *d* = 3.3), providing evidence that vision cues are taken into account in the combination condition for position, distance, and direction estimation. Observed variance in the combination condition does not differ significantly from the predicted integration but from the predicted alternation variance (Fig. 5), indicating that participants more likely combine vision and body cues than alternating between them.

Comparing the 0 tree and 400 tree environment, there is no significant difference consistent across individuals in weighting of visual and body cues between the environments but only a trend for higher vision weights in the 400 tree environment (Fig. 7A inserts). Observed weights in the 0 trees environment correlate with observed weights in the 400 trees environment for 2D Cartesian, distance, and direction estimation (Fig. 7A), indicating consistent cue weighting across environments.

Since cue weights are calculated from response variance and do not incorporate correctness of the final estimate it is interesting to understand if stronger weighting of vision cues also corresponds to better performance in the vision condition, or vice versa for body cues. Correlations of the vision weight with accuracy measured by position, direction, and distance error reveal that vision weight only correlates with position error in the vision condition–an effect that does not transfer to the other conditions (Tab. 1, supplement ‘**??**’). This indicates that participants tend to assign cue weights based on the consistency in their responses, independent of the correctness derived from the cue, or even if that cue systematically misinforms their final response.

**Table 1:** Correlations between observed cue weight and performance errors in each sensory condition. Each model,s vision cue weight is correlated with the performance error specific to its domain−2D Cartesian and position error, polar distance and distance error, polar direction and direction error. Correlations are described by their regression equation (y), p-value (p), and coefficient of determination (r2). Significance code: ∗ : p < 0.05, ∗∗ : p < 0.01, ∗ ∗ ∗ : p < 0.001. Regression plots are displayed in supplement, **??**,.

|  | observed vision weight: |  |  |
| --- | --- | --- | --- |
|  | 2D Cartesian<br>vs position error [m] | distance<br>vs distance error [m] | direction<br>vs direction error [°] |
| body<br>condition | $y = 8.2x + 9.0$ | $y = -3.7 - 2.8$ | $y = 43.5x + 11.8$ |
| | $p = 0.089$ | $p = 0.496$ | $p = 0.275$ |
| | $r^2 = 24\%$ | $r^2 = 4.3\%$ | $r^2 = 10.7\%$ |
| vision<br>condition | $y = -51.1x + 43.2$ | $y = -18.1 + 13$ | $y = -63.8x + 50.2$ |
| | $p = 0.001 **$ | $p = 0.115$ | $p = 0.328$ |
| | $r^2 = 63.5\%$ | $r^2 = 21\%$ | $r^2 = 8.7\%$ |
| combination<br>condition | $y = -10.5x + 14.4$ | $y = 2.7x - 2.6$ | $y = -39.6x + 35.7$ |
| | $p = 0.139$ | $p = 0.678$ | $p = 0.378$ |
| | $r^2 = 18.8\%$ | $r^2 = 1.6\%$ | $r^2 = 7.1\%$ |

In summary, cue combination of body and vision cues within path integration is considered unlikely to follow Bayesian cue integration principles because (1) we do not observe a reduction in response variance in the combination condition relative to either single-cue condition, and (2) observed cue weights do not correlate significantly with the predicted cue weights. Observed cue weighting appears consistent across environments with 0 or 400 trees, but generally does not correlate with task performance. Next, we analyse whether cue weighting of body and vision affects symmetric or asymmetric navigation behaviour between left and right-hand triangles.

### 2.3 Symmetric and asymmetric errors are affected by single-cue misestimations

To understand how individual preferences for body or vision cues affect navigation performance, we analyse symmetric and asymmetric error components, which we previously showed to be time-persistent for individual participants in triangle completion tasks [46]. We calculate symmetric and asymmetric direction and distance error components like in our previous study [46]. Symmetric direction and distance error components indicate the degree to under- or overshoot the target angle or distance (positive: overshoot; negative: undershoot), whereas asymmetric error components indicate directional biases to either side (from a participant’s ego-perspective: positive: right; negative: left). Since both, direction and distance error, are not significantly affected by the number of trees in the environment (Fig. 4) for the calculation of symmetric and asymmetric error components we pool the 0 and 400 tree environments (Fig. 8).

**Figure 8.**
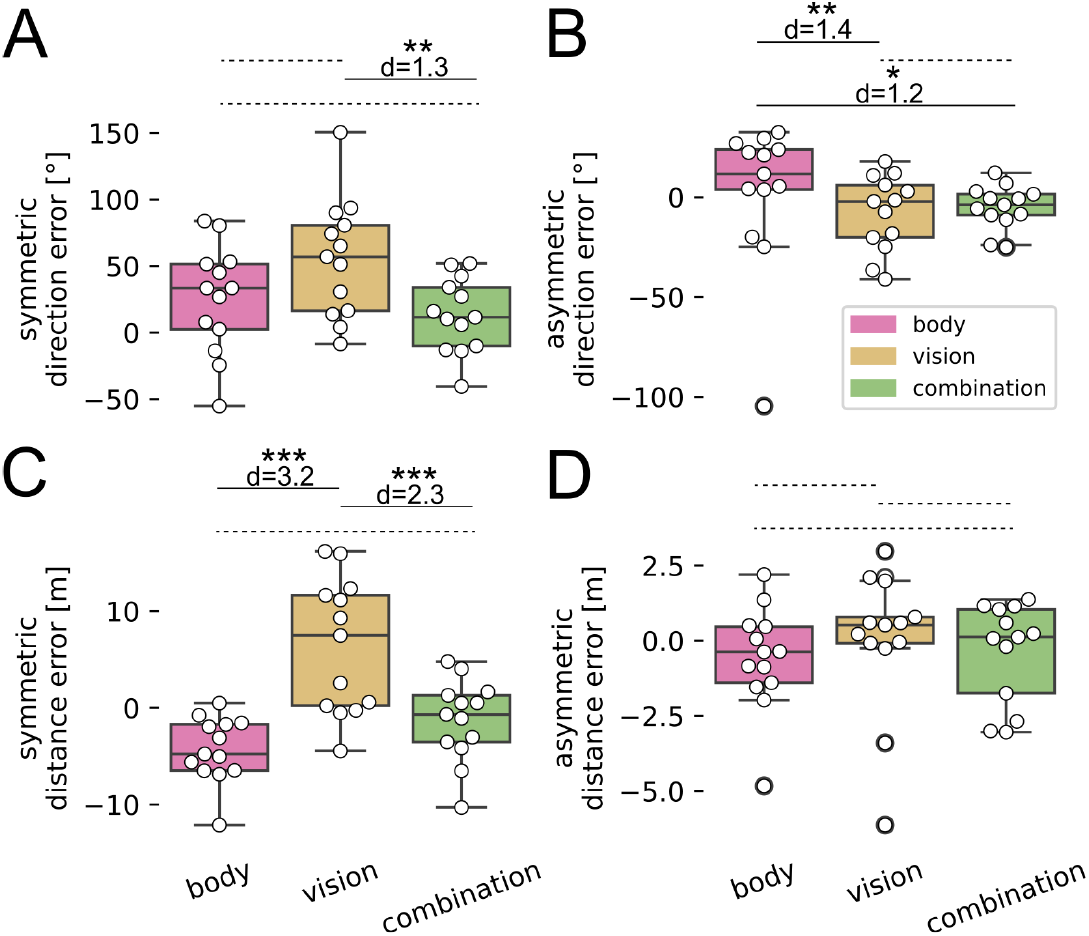
Symmetric (A, C) and asymmetric (B, D), direction (A, B) and distance error components (C, D) differ significantly between body, vision, and combination condition. Responses for trials with 0 or 400 trees are pooled. Data points represent single partici- pants’ means. Significance code (: p < 0.05, : p < 0.01, : p < 0.001) is based on Bonferroni corrected contrasts of estimated marginal means of LMMs using ’emmeans’ package (1.11.1) in R (4.5.0). Effect sizes are indicated by Cohen’s d.

Both symmetric and asymmetric direction errors show large inter-individual variability. While some participants systematically overshoot the correct target angle, others undershoot it (Fig. 8A). This is the case in both the combination and body conditions but not in the vision condition where the majority of participants overshoots the target angle by 55.3° on average (Fig. 8A). The symmetric direction error component is significantly larger in the vision condition (*mean* = 55.3°) than in the combination condition (13.3°, *d* = 1.3). The asymmetric direction error component is higher in the body condition (11.4°) than in the vision (−7.8°, *d* = 1.4) or combination condition (−4.7°, *d* = 1.2)(Fig. 8B). There is also large inter-individual variance in the distance domain. Participants on average overshoot the correct target distance in the vision condition (6.3 *m*) and undershoot it in the body condition (−4.3 *m*)(Fig. 8C). The symmetric distance error component in the vision condition (6.3 *m*) significantly differs from the body (−4.3 *m, d* = 3.2) and from the combination condition (−1.3 *m, d* = 2.3) which is–as we have seen above (section ‘Performance errors are larger for vision than for body condition’)–also reflected in differences of the vision condition in distance error (Fig. 4) and trajectory length (Fig. 3). Asymmetric distance error sizes are small and do not vary between conditions.

To understand the impact of individual symmetric and asymmetric biases in single-cue conditions on the naturalistic cue combination condition we correlate biases in the vision and body condition with the respective bias in the combination condition (Tab. 2, supplement ‘**??**’). Symmetric direction error in the combination condition significantly correlates with symmetric direction error in the body condition, but not with the vision condition (Tab. 2, 1st column). The asymmetric direction error in the combination condition, however, significantly correlates with the vision condition, but not with the body condition (Tab. 2, 2nd column). This might indicate that in triangle completion tasks symmetric over- or underestimation of angles (symmetric error) is caused by misestimation of body cues rather than of vision cues, and vice versa that asymmetric differences in angle estimation between left-hand and right-hand triangles might be caused by misestimation of vision cues rather than of body cues. The symmetric distance error in the combination condition significantly correlates with both, the body and the vision condition (Tab. 2, 3rd column), and even stronger with the sum of the symmetric distance error of both single-cue conditions (*y* = 0.4x − 2.1, *p* < 0.001, *r*^2^ = 73.9%). This might indicate that distance estimation in triangle combination tasks is affected by vision and body cues to the same degree.

**Table 2:** Correlations between each performance bias (symmetric, or asymmetric direction, or distance error) in single-cue conditions (body or vision) with respective bias in combination condition. These correlations indicate which biases observed in single-cue conditions are preserved in the combination condition. Correlations are described by their regression equation (y), p-value (p), and coefficient of determination (r2). Significance code: ∗: p < 0.05, ∗∗: p < 0.01, ∗ ∗ ∗: p < 0.001. Regression plots are displayed in supplement, **??**,.

|  | single-cue vs combination condition: |  |  |  |
| --- | --- | --- | --- | --- |
|  | symmetric<br>direction error[°] | asymmetric<br>direction error [°] | symmetric<br>distance error [m] | asymmetric<br>distance error[m] |
| body<br>condition | $y=0.53x-0.02$ | $y=0.08x-5.2$ | $y=0.95x+2.8$ | $y=0.28x-0.21$ |
| | $p=0.002 **$ | $p=0.376$ | $p=0.003 **$ | $p=0.324$ |
| | $r^2=60.5\%$ | $r^2=7.2\%$ | $r^2=57.4\%$ | $r^2=8.9\%$ |
| vision<br>condition | $y=0.29x-2.6$ | $y=0.46x-1.4$ | $y=0.46x-4.2$ | $y=0.24x-0.38$ |
| | $p=0.117$ | $p<0.001 ***$ | $p=0.003 **$ | $p=0.258$ |
| | $r^2=20.8\%$ | $r^2=64.3\%$ | $r^2=57.4\%$ | $r^2=11.5\%$ |

In summary, while participants in the vision condition mostly overshoot the correct target angle and distance, there is considerable inter-individual variance in the symmetric error component in the body and combination conditions. Observed weighting of vision and body cues correlates with symmetric direction and distance error components (high weight = low symmetric error), but not with asymmetric direction and distance error components (supplement ‘**??**’).

### 2.4 Gaming experience has no influence on performance

Since our experiment is essentially a VR video game we assessed gaming experience (hours per week during most active time on a 5-level grading scale: levels 1-5 ∼ 0, 1-2, 2-4, 4-6, >6) of our participants and correlate gaming experience with position error and cue weighting to understand if previous exposure to video gaming has a decisive impact on the interpretation of our study. We find large spread between participants in gaming experience in games in 2D (mean*±*sd: 2.7 *±* 1.7), 3D (2.3 *±* 1.8), with first-person perspective (2 *±* 1.5), third-person perspective (1.8 *±* 1.4), minimap (2.1 *±* 1.7), on PC (2.7 *±* 1.7), on console (2.1 *±* 1.4), on handhelds (2 *±* 1.5), or with VR headset (1.2 *±* 0.6). Of all gaming experience categories and their average, none significantly correlates with performance position error nor with vision cue weight in our experiment (supplement ‘**??**’). There is a tendency for participants to weight vision cues more when being more experienced in games with first-person perspective or minimap, but no effect to perform better. This holds true also when only considering performance in the joystick-controlled vision condition indicating that gaming experience did not influence performance, and that the controls in all conditions were manageable for our participants.

## 3 Discussion

This study assessed how participants weighted body and vision cues in the path integration process, using a naturalistic virtual reality (VR) triangle completion task. To the best of our knowledge, this is the largest-environment free-walking path integration study yet, conducted in a 1, 250 *m*^2^ gym. We provide a naturalistic yet controlled state of the art VR experience with no abandoned cues compared to natural navigation tasks. Previous studies have either limited vision information[14, 20], or body information[16, 21], or were conducted in comparably small space[4, 6, 9, 13, 18, 58, 59]. First, our experiment visually provides participants with rich ground texture and naturalistic trees for optic flow estimation while allowing free head movement. Second, concerning body cues the setup also allows to use all body cues that are known to inform navigation including vestibular cues from head turns and accelerations, proprioceptive cues from body position, efferent copies from limb movements, and somatosensory information from freely moving feet[20, 60, 61, 62].

### 3.1 Methodological considerations of isolating body and vision cues in path integration

This study focuses on the combination of vision and body cues within the path integration process. Path integration describes the continuous tracking of the walked path and is informed during the outbound path and retrieved during homing. The outbound path can be referred to as the encoding phase and homing as the execution phase, where the encoding process continues to influence execution[35]. Since the path integration process takes place during the outbound and homing phase we ensure in our single-cue trials to limit information to only vision or only body cues during the whole trial including the outbound and homing phase. In previous research, Nardini et al. [14], Zhao and Warren [4], and Chen et al. [18] (upon others) calculated cue weighting of vision and body cues by separate single-cue and combination trials that all provided both, vision and body cues, during the outbound path allowing encoding by both cues. Only at the end of the outbound path, before homing, the world was rendered invisible for body trials, and for vision trials participants were disoriented on a swivel chair. Swivel chairs disoriented participants on the spot essentially disturbing the direction component of the path integrator but leaving the distance component intact. The intact distance estimate of the path integrator, that was informed also by body cues during the outbound path, most likely reduced variance in response distance and thus, also any 2D variance measure or subsequent cue weighting calculations. Hence, vision-only trials might be influenced by body cues in some of the previous research[4, 18]. This difference to our study is important to understand because the focus of previous studies was cue weighting in the execution of targeted homing including path integration and landmark guidance, whereas our focus is on cue weighting within the full process of informing and retrieving the path integration system without the incorporation of landmark guidance.

Kearns et al. [6] also explored the integration of visual and body-based cues in path integration. However, their body-only condition still relied on visual posts for guidance. In addition, the used VR headset had a more restricted field of view (horizontal: 60° vs. 110°; vertical: 40° vs. 96°), and the experimental arena was smaller (144 *m*^2^ vs.1, 215 *m*^2^). While comparable to our study, these limitations retained residual optic flow and constrained visual perception, which our setup was designed to minimise.

However, strictly separating single-cue conditions throughout the outbound and homing path in this study necessarily induced distinct differences between the conditions. In body trials, participants are challenged with following the controller vibration guidance, and in vision trials participants need to translate and rotate their avatar with joysticks. Differences between the conditions including longer trajectories and higher speed in vision trials and lower speed in body trials (Fig. 3) cannot clearly be ascribed to differences in sensory processing but need to incorporate the aforementioned methodological differences in trial execution. However, the fact that gaming experience does not correlate with task performance provides good evidence that all conditions can be executed also by gaming-naive participants. Yet another methodological limitation of our study is the physical limitation of the space. We conducted the yet largest-environment free-walking VR navigation study in a 45 *m ×* 25 *m* gym and wanted to make the triangle completion task as large as reasonably possible. Although we arranged the triangles in a way that leaves much room for misestimations in distance and direction relative to the target (Fig. 9), participants walked out-of-bounds in about every fourth trial in the body and combination condition (Fig. 2). Since participants are stopped before walking into walls, implicit learning in form of ‘this was not the right way’ could not be controlled for. Additionally, some participants might have been afraid to walk into a wall and might have favoured shorter paths than in real-world environments. Such fear might also have affected straightness, trajectory length, walking speed (Fig. 3), and distance estimation (Fig. 4) in the blindfolded body condition in which some participants reported initial discomfort. Eventually discomfort may also have influenced the cue weights derived from navigation performance. Future studies would benefit from an open space that avoids unwanted feedback or discomfort for participants.

**Figure 9.**
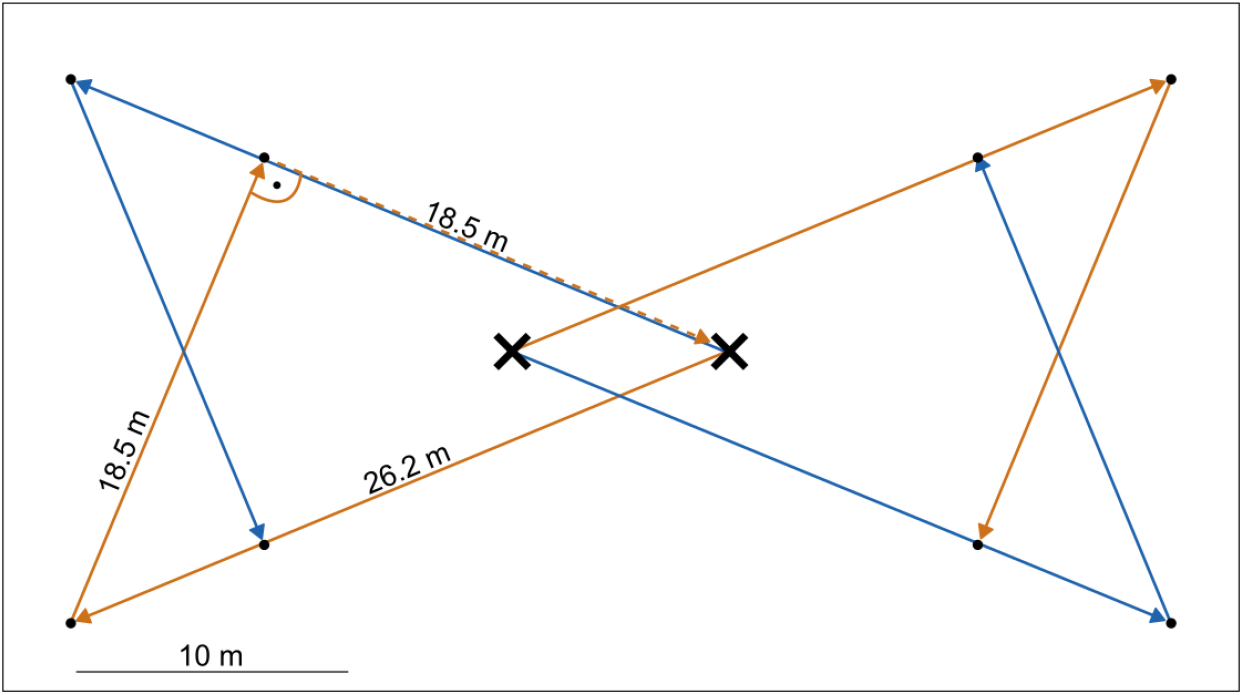
Placement of waypoints (black) for triangle completion task in the gym. Right-hand triangles (orange) and left-hand triangles (blue) were placed in all four corners of the gym. All triangles are isosceles and share the same dimensions with requiring participants to turn 90° left or right and walk straight for 18.5 m to ideally complete the task.

### 3.2 Body cues dominate vision cues in non-Bayesian homing strategies

Observed cue weights, calculated from relative nearness of combination responses to either single-cue response, do not correlate with predictions from the Bayesian cue combination models, calculated from relative variance in either single-cue condition (Fig. 7B). This means that participants in our experiment do not integrate vision and body cues following optimal Bayesian cue combination principles, in neither 2D, distance, nor direction domain, neither without trees nor with 400 trees in the environment. As pointed out by Scheller and Nardini [63], it is essential to test for optimal cue integration separately for each participant rather than across the population. However, also when considering the sensory condition with the lowest response variance for each participant, we find no evidence of variance reduction in the combination condition relative to the single-cue condition for 2D Cartesian, distance, or direction estimation (Fig. 5). These results further suggest that the combination of vision and body cues in the path integration process does not follow Bayesian cue integration principles.

Unlike other studies[4, 14, 18, 28] who reported their participants to integrate selfmotion path integration cues and visual landmark cues near-optimally in the execution of homing, we do not find evidence for this form of cue combination for body and vision cues within the path integration system. In line with this, recent research on test–retest reliability has shown low intra-class correlations for precision and cue weighting in a triangle completion paradigm[64]. This aligns with recent findings in *Drosophila*, where individual differences in the trade-off between stability and flexibility of spatial representations argue against the assumption of consistently optimal navigation [65]. Additionally, cue alternation–alike observed for the combination of landmarks and path integration in children in the past[14]–is considered unlikely for our participants as we find significant differences between observed and predicted response variance for the alternation model (Fig. 5). Since prior beliefs can strongly affect cue integration[19, 22, 27], the observed dominance of body cues in our study might also stem e.g. from a previously acquired distrust in the visual world presented on the VR headset.

How might vision and body cues be integrated in our experimental paradigm if not according to Bayesian cue integration principles? Response variance in the combination condition does not differ from the predictions of the cue integration models but also not from the body condition (Fig. 5). Hence, it might be that response precision (variance) is dominated by body cues as it proves to be the more reliable cue for most participants (Fig. 7). Possibly, it might also be that response variance in the body condition was already too low to reduce it even further with additional vision cues in the combination condition. Response accuracy, however, follows the direction of body and vision cues (Fig. 4C), and falls between the single-cue estimates for distance estimation (Fig. 4B). These findings indicate that path integration in our VR task is primarily governed by body cues, with vision contributing only to distance estimation.

Knowing that Bayesian cue combination is unlikely to play a role for the path integration process investigated in this study, we can still use observed cue weights to get an understanding of the importance of body and vision information. Position and distance errors are higher in the vision condition which cannot be explained by gaming experience. Instead it appears that visual cues are assessed in a way that eventually makes them less reliable for participants. This agrees with the findings by Chance et al. [13] who observed worse performance in a virtual maze task when participants used only vision cues with joystick controls than when participants actively walked. In the vision condition, distance errors are higher than in the other sensory conditions, and direction errors are similar. Since distance is assessed by self-translation this indicates that translational optic flow is assessed differently in vision trials (Fig. 4A and B). However, as direction is assessed by self-rotation, rotational optic flow appears to be assessed similarly with body cues, vision cues, and both (Fig. 4). These findings support Kearns et al. [6] who suggest that translational and rotational optic flow differently affect homing performance and in conjunction might indicate that homing distance and homing direction are estimated separately in triangle completion tasks. In contrast, they found that participants switched from underturning the target angles with joystick control to overturning when actively walking. This effect is not supported by our study and might be caused by rather unnaturally low rotation speed with joystick controls in their study (30°/*s* vs 60°/*s* in our study).

Our findings, together with those of Kearns et al. [6], propose that homing direction and distance are processed separately and support the rationale for contrasting cue combination in Cartesian versus polar space. Conceptually, the Cartesian model assumes that participants integrate all sensory information to estimate a single 2D endpoint position, akin to locating themselves on a map. In contrast, the polar model reflects a process in which distance and direction are computed and combined separately, aligning more closely with the above indications.

In the path integration process, we find body cues to be more reliable than visual cues and correspondingly weighting of body cues to dominate (Fig. 7). This contrasts with Ehinger et al. [9], who reported no behavioural differences when manipulating proprioceptive and vestibular cues in a triangle completion task what might be a consequence of their rather small sample size (*N* = 5). Our findings, however, align with Kearns et al. [6], who found body cues to dominate in triangle completion, and with Adamo et al. [66], who reported superior navigation with vestibular cues in a wheelchair compared to visual cues alone. Further support for the importance of body cues comes from Xie et al. [67], who showed that patients with vestibular loss made larger distance and angle errors than healthy controls in a triangle completion task. Together, these findings underscore the critical role of body cues including vestibular, proprioceptive, and efferent signals which also proved to be more reliable than visual information in our experiment (Fig. 5).

Nevertheless, visual cues are taken into account in path integration, as indicated by mean vision weights being significantly greater than zero (95% − *ci* : [0.2, 0.31]). Consistently, Tcheang et al. [68] demonstrated in a triangle completion task that vision and body cues were integrated into a unified multimodal representation, rather than independently influencing action. Our results support this view, suggesting that vision and body cues are fused into a single path integration estimate.

### 3.3 Vision cues might cause overturning - body cues might cause side biases

Independent of the origin of differences between the conditions they naturally also manifest themselves in calculated individual biases. For the vision condition, symmetric direction and distance errors for the majority of participants are positive (Fig. 8A and B) meaning they overshoot the target distance and angle (Fig. 4B and C). In the body condition, underestimation of the target distance (Fig. 4B) implies negative symmetric distance errors (Fig. 8C). Since in distance estimation symmetric over- or undershoots are best explained by the sum of misestimating the target distance in the body and the vision condition (Tab. 2), it seems that overshooting the target distance with only vision cues can be compensated when undershooting with only body cues (Fig. 8C). This, however, does not seem to be the case for direction estimation where symmetric error in the combination condition only correlates with the body condition and asymmetric error in the combination condition only correlates with the vision condition (Tab. 2). Hence, the general tendency for overturning the target angle in our experiment (positive symmetric direction error) might stem from body cues, whereas, individual side biases (asymmetric direction error) might rather originate from misestimations of vision cues (Tab. 2). Overturning the target angle was interpreted as a functionally useful behaviour in desert ants because they cause the return path to cross the first leg of the outbound journey, enhancing the chance to identify salient landmarks [69] which, however, were absent in our experiment. The latter observation that individual side biases stem rather from misestimations in the encoding of optic flow supports our hypothesis proposed in a previous study, as no body cues were available in that study [46]. In contrast, Jetzschke et al. [56] attributed directional biases in rotation estimation to vestibular processing, as participants in their study were blindfolded.

Since asymmetric direction error components do not correlate with cue weighting, there is no evidence that cue weighting affects side biases nor that participants adapt their cue weighting relative to their side bias (supplement ‘**??**’). Our initial expectation that participants who weight body cues stronger than vision cues have smaller side biases is not supported by the data. Symmetric error components, however, correlate well with cue weighting with strong weighting of vision cues predicting low symmetric error components in the vision condition, and high ones in the body condition. It is unclear whether cue weighting affects symmetric error components or vice versa.

### 3.4 Ambiguous forests and fallow land do not influence cue weighting or navigation performance

There is no significant effect of weighting vision cues more in the 400 tree condition (Fig. 7A inserts). We observe a consistent cue weighting strategy across the 0 and 400 tree environment for 2D position, distance, and direction. This corresponds to Kearns et al. [6] who find that when participants actively walked, there was no behavioural difference with or without texture on the VR wall and floor. Direction estimates in our experiment, however, are significantly less variable with 400 trees (Fig. 5, supplement ‘**??**’ direction variance). In contrast, Chen et al. [18] observed that participants assigned larger vision weights in a ‘rich’ environment with three distinct landmarks than in a ‘poor’ environment with only one post. Crucially, their environments manipulated the number of unambiguous landmarks for landmark guidance, rather than focusing on the amount of optic flow for path integration as in our study. In our setup with numerous ambiguous trees, we cannot validate an effect of environment on weighting of body and vision cues for fallow land versus a forest. While richer 3D texture and increased optic flow could be expected to facilitate accurate translation and rotation estimation (Fig. 4) we do not observe participants adapting their vision weights in the forest. Salient landmarks seem to be necessary to distinctly affect weighting of visual cues and error reduction in navigation. Notably, one needs to incorporate that the environment with 400 trees potentially allows participants besides path integration to use landmark guidance. This means that the cue combination process we observe is not limited to the path integration system in the 400 tree environment. However, the trees are ambiguous and provide no clear advantage for participants in returning to their home location, as reflected by similar performance errors (Fig. 4).

### 3.5 Redundancy and sensory specialisation of landmark guidance and path integration

While landmark guidance relies on visual cues, our findings suggest that path integration is primarily driven by body-based information. In principle, both systems support goal-directed navigation[4, 18, 28, 59], indicating a synergistic rather than a hierarchical relationship. From an evolutionary perspective, their dependence on distinct sensory modalities likely reflects an adaptive strategy that promotes redundancy and resilience: if one sensory channel is compromised, the other can still support navigation towards essential resources or shelter. In accordance with this, Zanchi et al [70] investigated the integration of visual and auditory landmark cues and found that visual cues generally dominated landmark guidance [70]. Cue integration consistent with the Bayesian framework was only observed for those participants who weighted visual and auditory cues similarly. In contrast, individuals who showed a strong preference for visual cues did not integrate the cues in a Bayesian manner. In the present study, we did not observe such distinct subgroups with divergent cue weighting patterns, supported by the narrow 95%-ci of the mean vision weight across participants(2D Cartesian: [0.21, 0.31], distance: [0.19, 0.31], direction: [0.35, 0.48]).

Taken together, the findings from Zanchi et al. [70] and the present study suggest that landmark guidance is largely driven by visual input, and that path integration is dominated by body-based cues, with no consistent evidence for Bayesian cue integration within either system individually. Instead, Bayesian integration appears to occur only after both the landmark estimate and the path integration estimate have been consolidated independently[4, 18, 25].

### Conclusion

This study provides evidence that, in the path integration system most participants substantially favour body cues over vision cues in a non-Bayesian combination of both cues in our naturalistic large-environment VR task. There is considerable inter-individual variance in the strength of this cue dominance. Strategic weighting of body and vision information appears constant across a sparse and a cluttered environment. Going from fallow land to a forest reduces directional variance what, however, does not make participants assign significantly higher weights to visual cues. We found no evidence that participants’ side biases were directly connected to cue weighting.

## 5 Acknowledgment

This manuscript has been submitted as one chapter in the dissertation titled “Mechanisms of Multi-Guidance in Human Navigation” by J.S. at Bielefeld University and will appear online within the next month.

## 6 Data Availability

The datasets generated and analysed during the current study are available in Bielefeld University’s PUB repository at https://doi.org/10.4119/unibi/3011122. Analysis code with accompanying instructions for execution is available on GitLab at https://gitlab.ub.uni-bielefeld.de/jonas.scherer/embodied-navigation-whole-body-movement-drives-path-integration-in-large-scale-free-walking-virtual-reality Implementation details of the Unity3D project and an executable Unity project of this study’s task are available on GitLab at https://gitlab.ub.uni-bielefeld.de/virtual_navigation_tools/unity_vnt_showcase_triangle_completion.

## 7 Contributions

Conceptualisation: J.S., A.K., and N.B.; Methodology: J.S.; Software: J.S.; Investigation: J.S., A.K.; Data curation: J.S., A.K.; Visualisation: J.S.; Formal Analysis: J.S.; Validation: A.K.; Writing-original draft: J.S; Writing-review & editing: J.S., A.K., M.E., N.B.; Funding acquisition, project administration, resources, and supervision: M.E., N.B.

## 9 Methods

### 9.1 Experimental design

This study explores how idiothetic body (vestibular and proprioceptive) cues and visual (optic flow) cues are combined in the path integration process of individuals and how this combination is affected by different environments. For that, we analyse behavioural performance of individuals in a large-environment VR free-walking triangle completion task in three different conditions, each with and without a tree clutter. We manipulate participants’ accessibility to body cues and visual cues in a vision condition with only visual cues, a body condition with only body cues, and a combination condition with both modalities of cues.

### 9.5 Experimental procedure

*N* = 13 healthy participants (aged 32.2 *±* 12.2 (mean*±*sd) years, 8 self-identified as female, 5 as male) took part in the experiment. Participants were informed about the general procedure of the experiment but not about its specific purpose. All participants gave their written informed consent to participate and received monetary compensation (10 Euros per hour). The experiments were approved by the Bielefeld University Ethics Committee and conducted in accordance with the guidelines of the Deutsche Gesellschaft für Psychologie e.V. (DGPs), which correspond to the guidelines of the American Psychological Association (APA). All data collected from the participants and during the experiment was handled according to the General Data Protection Regulation (GDPR) of the European Union.

The experiment was carried out over four sessions (1 training + 3 test sessions). It was created using the “Virtual Navigation Toolbox“[71] for the Unity3D game engine and was executed on the Meta Quest 3 head-mounted display (resolution: 2064*×*2208 per eye; field of view: 110° horizontal, 96° vertical) in the 45 *m×*25 *m* = 1, 215 *m*^2^ gym at Bielefeld University. The triangle completion task was presented in form of a VR task in which participants collect firewood and return to their campfire (see ‘Virtual reality triangle completion task’ for details). In the combination condition, participants actively walk around and have access to body cues and visual cues. In the body condition, participants actively walk around but are blindfolded by a uniform grey screen in the VR headset, denying visual cues. In the vision condition, participants stand still and use joysticks to move, denying body cues.

Each test session consisted of 20 pseudo-randomised trials. Trials in the combination and vision condition were each executed without tres and with 400 trees in the surround, each in 2 right and 2 left triangles, contributing 16 trials. The body condition was conducted in 2 right and 2 left triangles only without tree clutter because participants were blindfolded anyways, contributing 4 trials. Over three test sessions this yields *n* = 6 repetitions per condition (combination, vision, body + trees, no-trees) and triangle side (left or right) for each participant. Left and right mirror symmetric triangles were located in 4 locations within the gym with 2 different starting locations (Fig. 9). Testing the triangle on both sides is necessary to calculate the symmetric and the asymmetric individual error components [46](see ‘Error calculations’ for details). An experimenter instructed and assisted participants according to a standardised protocol during 6 initial training trials, with two trials in each sensory condition and smaller, differently shaped triangle paths to ensure task and controls to be well understood without anticipating the paths during testing. After training, participants completed the remaining test sessions without further assistance. Participants were instructed to take regular breaks in between experimental sessions of at least 5-10 minutes. Sessions on average took 27*±*10 (mean*±*sd) minutes to complete.

After the test sessions, participants completed questionnaires about their gaming experience and the strategies they employed during the experiment.

### 9.3 Virtual reality triangle completion task

We tested participants in a Meta Quest 3 VR large-environment triangle completion task designed with the “Virtual Navigation Toolbox”[71]. In our VR version of the triangle completion task, the avatar was controlled from a first-person view depending on the sensory condition of the trial. In the combination and body condition participants moved by active walking, and in the vision condition by using the left or right joystick, for translation and rotation, respectively. At the beginning of each trial participants were informed by audio and text instructions to move physically or to stand still and use the joysticks. When moving physically through the gym, participants got an audio and text alert when they approached walls closer than 1.5 *m*. For safety, an experimenter followed the participants to acoustically interfere when they were getting too close to the walls. When using joysticks for avatar movement, a yellow banner covered the field of view when participants moved more than 0.1 *m* or rotated more than 10° and reminded participants to stand still.

To complete the triangle completion task in the VR, the participants were first guided to their home location indicated by a stylised campfire. This initial approach started wherever the previous trial ended (Fig. 9). The campfire then disappeared and an arrow at the bottom of the screen guided the participant to collect firewood logs at two distinct locations, while being able to move freely. This was termed the outbound phase of the trial. To ensure an overall straight path along the legs of the outbound path, the participant was confronted with a time limit (100 *s*) to collect all the firewood. The remaining time was always visible in the form of a timeout bar at the top of the screen, also in the body condition where the screen was otherwise completely grey. In the body condition, participants were guided along the outbound path by vibrations of the handheld controllers. Adapted from Harootonian et al. [20], when participants deviated more than 10° from the correct direction towards the next waypoint, vibrations indicated the need to turn more towards the opposite side, and vibrations got stronger at deviations above 30°.

Firewood log waypoints were collected when participants got as close as 0.1 *m* in vision or combination trials, or as close as 0.4 *m* in body trials to ease trial execution. After collecting the second firewood log, participants were again presented with a 60 *s* timer, and had to find back to the campfire location (i.e. the goal location), entering and confirming their response via button press. Neither the firewood logs nor the campfire were visible during the return, to prevent their use as spatial cues. After the participant had indicated reaching the goal, the next trial was started by regenerating the environment according to the new trial’s sensory condition and environment. For forest environments with 400 trees, a tree clutter was randomly chosen from a set of five clutters for each respective trial. Tree clutters were generated by randomly placing 400 trees in a circular area with 100 *m* radius and manually repositioning trees that obstruct the straight path between experimental waypoints. No feedback on performance was given in training or test sessions. Audio instructions during the trial informed participants about what to do and whether to move physically or by using the joysticks. During each trial, the position and rotation of the player’s avatar in the virtual environment were recorded for every rendered frame. Identical grass tufts on the ground were randomly arranged for each trial to enable visual estimation of self-motion by optic flow. A sun-like light source provided global illumination at 90° azimuth. When moved by joysticks in the visual condition, the avatar moved at 2.5 *m*/*s* and turned at 60°/*s*. Virtual objects were scaled to realistic sizes (trees: 21 *m*, grass tufts: 0.3 *m*). The experiment rendered on the headset at an average framerate of 39 frames per second. Preliminary validation trials at multiple task-relevant distances showed tracking accuracy of the VR headset at single digit centimeter level. All headsets were cleaned with disinfectant wipes before use. Disposable hygiene covers were attached and sprayed with deodorant close to the nose, to cover any olfactory cues in the gym. During the experiment the headset played music chosen from a list of 10 royalty free songs by the participants, to cover any acoustic cues in the gym. Participants were free to walk on socks or bright-soled shoes.

### 9.4 Data analysis

#### 9.4.1 Error calculations

We calculated position errors, distance errors, and direction errors from homing response endpoints. Position error is the Euclidean distance from the endpoint to the target position. Distance error is the degree of walking too far (positive) or not far enough (negative). Direction error is the directional deviation from the target angle and defined positive for deviations to the right and negative for deviations to the left. Since in the VR triangle completion task waypoints were triggered at 0.1 *m* (vision and combination condition) or 0.4 *m* (body condition) physical distance, we used the real physical position of each trial’s homing start instead of the programmed waypoint location for error calculations. From the observed direction and distance errors in left and right-hand triangles, we calculated distributions of the symmetric and asymmetric error components at individual participant level as in one of our previous studies [46]. The degree of symmetric direction error component (*direrr*_*sym*_) was calculated by:

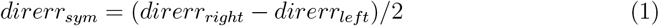

where *direrr*_*right*_ is one direction error in a right-hand triangle trial and *direrr*_*left*_ is one direction error in a left-hand triangle trial. This component is the shared degree of over- or underturning in left and right turns. The asymmetric direction error component (*direrr*_*asym*_) is calculated by:

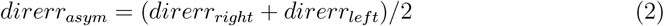

This component describes the tendency of a participant towards the left (negative sign) or right (positive sign) side. One can think of the asymmetric error component as the difference between the tendencies to over- or undershoot left or right turns. For analyses we then took the mean of the resulting distributions from all combinations of one left triangle response and one right triangle response.

Symmetric and asymmetric distance error components were calculated accordingly (see [46] for details). The symmetric distance error component describes the tendency of a participant to over- or undershoot the target distance symmetrically for left and righthand triangles. The asymmetric distance error component describes the tendency of a participant to over- or undershoot the target distance for one triangle side systematically more than for the other.

#### 9.4.2 Bayesian cue combination models

We modelled cue combination of visual and body cues in the path integration system as a Bayesian cue integration and as a cue alternation model. We calculated cue integration and cue alternation models in a Cartesian coordinate system alike Chen et al. [18], and additionally in a polar coordinate system. In both coordinate systems the origin was set to be the homing start location and the direction towards the homing target as the positive y-axis, or *θ* = 0 respectively. Based on the equations used for the Cartesian model[18] we will explain the polar model afterwards. Any deviations from Chen et al. [18] are pointed out below, where applicable.

#### Cartesian model

Following Bayesian cue integration principles, we modelled a cue’s weight as its reliability in this cue’s single-cue condition relative to the other cue’s single-cue reliability. A cue’s reliability (*r*) is the inverse of the observed variance (*σ*^2^) in the respective single-cue condition:

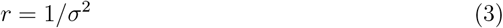

For the Cartesian models we followed the variance calculation described by Chen et al. [18], where observed variance is defined as:

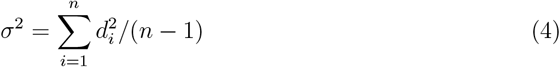

with *d* being the distance of a response from the centroid of its respective condition and *n* being the sample size within that condition. Following this, predicted vision weight was calculated by:

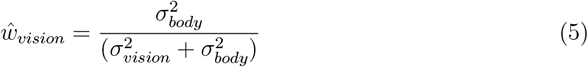

and, assuming absence of other cues in our experiment,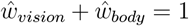. Correspondingly, predicted body weight was calculated by:

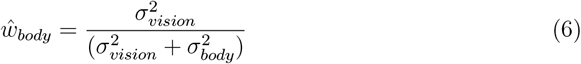

When cue integration in the combination condition follows Bayesian cue integration principles this predicts reduction of response variance in the combination condition upon the single-cue variances by:

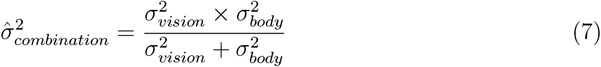

When participants, instead of integrating cues, alternated between them at the ratio of the single cue weights, combination variance was calculated by an alternation model. Here, our calculations deviated from Chen et al. [18], who use the one-dimensional x-coordinate as ‘a proxy for angular heading’, which is sensible in their comparably small triangles. For our large-scale triangles, we considered both x and y-coordinates equally in our Cartesian cue alternation model and extend their model by representing each singlecue condition as a bivariate Gaussian distribution defined by its mean *µ* and covariance matrix *cov*. The resulting mixture distribution’s mean is equal to the weighted average of the two single-cue centroids. The distribution’s covariance matrix is given by the sum of the weighted cue covariances and the covariance of the cue means. Specifically, the total covariance matrix was computed by:

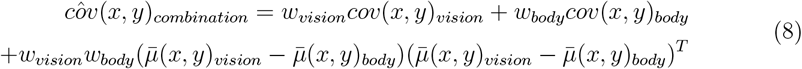

with *cov*(*x, y*) being two-dimensional covariance matrices, and 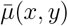 being mean responses. To express response variance in the same scalar form used for observed data (see equation 4), we computed the trace of the alternation model’s covariance matrix, which corresponds to the expected squared Euclidean distance of a sample from the centroid:

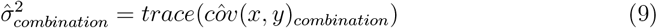

We could assess whether the cue combination process of vision and body cues in the path integration system follows Bayesian cue integration or alternation principles by comparing the predicted weights and variances with the observed ones. Observed cue weights were calculated by the relative proximity of responses in the combination condition relative to the responses in single-cue conditions by:

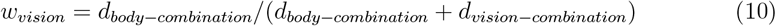

(and likewise for *w*_*body*_), with *d*_*body*−*combination*_ being the Euclidean distance between the response centroid in the body condition and the combination condition. Observed variance in the combination condition was calculated alike in single-cue conditions by equation 4. We also predicted mean response estimates, which we did not use for any analytical model assessments, but only for plotting (Fig. 6). Response estimates were predicted equally for integration and alternation models, because both are linear combinations proportional to cue weighting, by:

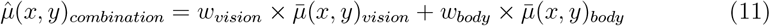

with 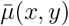 being the mean response in the respective condition.

### Polar model - distance component

In the polar models we calculated predicted variances and weights separately for the distance *ρ* and directional *θ* component (see also [16]).

The distance component was modelled as a one-dimensional Gaussian defined by mean *ρ* and standard deviation *σ*. Observed *σ* and *ρ* were fitted by maximum likelihood estimation (MLE) for each condition. From there, predicted weight calculations are equal to the Cartesian model (see equations 5 - 7). For cue integration, predicted response variance in the cue combination condition also corresponds to the Cartesian model (see equation 7). For cue alternation, however, the polar distance component is a one-dimensional measure, and hence, we use the same equation as Chen et al. [18] (equation 14):

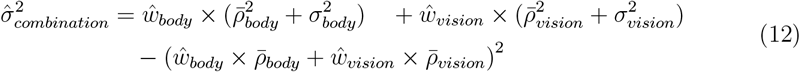

Observed weights were calculated like in equation 10, but with the absolute 1D difference between mean distance estimates instead of the Euclidean distance between response centroids. We computed the predicted response distance for the integration and alternation polar model also just for plotting (Fig. 6), without considering it for any analytical model assessments, by equation 11 with one-dimensional 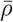 instead of 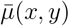.

#### Polar model - direction component

The direction component was modelled as a von Mises distribution (circular normal distribution) defined by mean direction *θ* and concentration coefficient *κ* (as a rule of thumb: *κ* ≈ 1/*circular sd*)[4, 57]. Observed *κ* and *θ* were fitted by MLE for each condition[72]. Since *κ* is directly proportional to a cue’s weight (not the inverse as for variance) the predicted vision and body weights were calculated by:

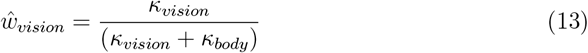

and

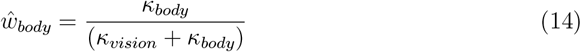

respectively. Following Murray and Morgenstern [57] (equation 15), predicted concentration for cue integration in the combination condition was calculated by:

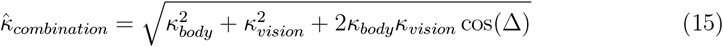

with Δ = *θ*_*body*_ − *θ*_*vision*_.

For predicted concentration with cue alternation, we approximated the probability-based mixture of the two single-cue von Mises distributions by a single von Mises with the same first circular moment[73]. Specifically, we computed the mean resultant length *R* as the magnitude of the weighted mixture vector 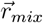 formed from the mean directions and mean resultant lengths of the single-cue conditions:

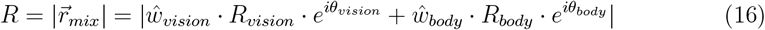

The concentration κ was then obtained by numerically inverting the relationship *R* = *I*_1_(*κ*)/*I*_0_(*κ*) with *I*_0_ and *I*_1_ being the modified Bessel function of order 0 and 1, respectively[73, 74].

Observed weights were calculated like in equation 10, but with the absolute circular difference between mean direction estimates instead of the Euclidean distance between response centroids. We computed the predicted mean direction response for cue integration and alternation but did not use it for any analytical model assessment but only for plotting (Fig. 6). Since we observed substantial differences between response directions in the vision and body condition, we used the equation in Zhao and Warren [4](supplement equation 1) that considers a directional difference in single-cue responses. Predicted mean direction response in the cue integration model was calculated by:

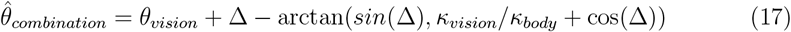

with *arctan* being the four-quadrant inverse tangent and again Δ = *θ*_*body*_ − *θ*_*vision*_. Predicted mean direction response in the cue alternation model is given by the direction of the weighted mixture vector 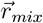 in equation 16.

For direct comparison with the other models’ standard deviations we transformed concentration *κ* into circular sd following Frischkorn and Popov [75]:

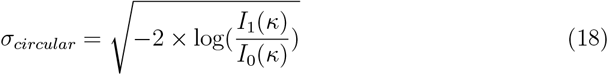

#### Pooling of trials

Bayesian models, were computed for a standardised right-hand triangle only. Due to rather small sample size per triangle side in each condition (*n* = 6) we instead mirrored responses from left-hand triangles at the y-axis and pooled responses from both sides. To not artificially increase variances, we moved endpoints of each triangle side so that its respective mean measure (Cartesian: centroid, distance: mean distance, direction: mean direction) aligns with the origin, or *θ* = 0 for von Mises, before pooling and fitting the variance parameter. Observed and predicted mean responses (*µ*, or *θ*) and observed weights (*w*_*vision*_ and *w*_*body*_) were calculated separately for left and right-hand triangles and averaged afterwards.

### 9.5 Data processing

Walked triangles from both home locations within the gym (Fig. 9) were pooled. We standardised all trials to the same coordinate system so that the target location was at the origin and the correct homing direction was aligned with the y-axis. We excluded trials in which participants walked out of bounds during the outbound path and did not collect all waypoint firewood logs. We excluded outliers in the same way as Chen et al. [18]: Responses that exceeded the third quartile by more than three times the interquartile range (IQR) in the distance to their respective centroid were excluded, where the IQR was calculated from pooled endpoints of right and left-hand triangles. For outlier detection we compared each endpoint to its respective centroid (each combination of vision – body - combination, left - right hand triangle, and 0 - 400 trees was checked separately). On average 2.7 trials per participant were excluded from analysis.

### 9.6 Power analysis

As a compromise between response variance in our previous study on individual persistence of symmetric and asymmetric error components (see [46]) and a realistic experiment duration, we decided to conduct *n* = 6 repetitions per participant (*N* = 13), per condition (vision, body, combination), per environment (0 or 400 trees, except for the body condition because participants were blindfolded anyways), and per triangle side (left or right). Based on the condition with highest mean variance across participants for each domain (Cartesian 2D, distance, direction) we quantified power of our study by simulating data from a distribution with the respective variance with *n* = 12 samples. We used *n* = 12 samples and not *n* = 6 samples because in our analysis we either pooled across triangle sides for performance and cue weighting calculations, or across number of trees for bias calculations. For direction, we observed the highest variance in the vision condition with 0 trees at *κ* = 6.1^° −1^, at which sample size in our study can resolve circular sd of 7° at 80% power. For distance, we observed the highest variance in the vision condition with 400 trees at *sd* = 6.4 *m*, at which sample size in our study can resolve distance sd of 1.8 *m* at 80% power. For Cartesian 2D estimation, we observed the highest variance in the vision condition with 0 trees at *sd* = 15.3 *m*, at which sample size in our study can resolve Cartesian 2D sd (equation 4) of 4.3 *m* at 80% power. Power in the body and combination condition are considerably higher (direction: 4.7°, distance: 1.0 *m*, Cartesian 2D: 1.8 *m* at 80% power).

### 9.7 LLM usage

We utilised GPT-5 from OpenAI during the writing and programming procedure of this manuscript.

## 10. Figures

