## Supplementary Material for "Embodied Navigation: whole-body movement drives path integration in large-scale free-walking virtual reality"

#### S1 Linear mixed models

Position, distance, or direction error and as well the respective variances in the error measures are predicted from the fixed categorical effects 'condition' and 'number of trees' and a random 'participant' effect (error $\sim$ number of trees + condition + (1|participant)). To assess the effect of number of trees we remove the body condition, which does not have a variant with 400 trees, allowing to also include an interaction between the fixed effects (error $\sim$ number of trees $*$ number of trees + (1|participant)). Below are the detailed results of all linear mixed models. Model results were exported from R using the 'texreg' package[1]. 'w/o' stands for 'without'.

**position error**

Table 1

| position error ~ number of trees + condition + (1\|participant) | | | | | | |
|---|---|---|---|---|---|---|
| | Sum Sq | Mean Sq | Df1 | Df2 | F | p |
| c(number of trees) | 1725.4 | 1725.4 | 1 | 723.10 | 18.922 | 1.558e-05 *** |
| c(condition) | 22850.9 | 11425.5 | 2 | 723.07 | 125.295 | < 2.2e-16 *** |
| AIC | 5467.45 | | | | | |
| BIC | 5495.08 | | | | | |
| Log Likelihood | −2727.73 | | | | | |
| Num. obs. | 739 | | | | | |
| Num. groups: ID | 13 | | | | | |
| Var: ID (Intercept) | 20.15 | | | | | |
| Var: Residual | 91.19 | | | | | |

$^{***}p < 0.001$; $^{**}p < 0.01$; $^{*}p < 0.05$

Table 2

| position error ~ number of trees ∗ condition + (1\|participant) (w/o body condition) | | | | | | |
|---|---|---|---|---|---|---|
| | Sum Sq | Mean Sq | Df1 | Df2 | F | p |
| c(number of trees) | 1694.5 | 1694.5 | 1 | 589.08 | 18.125 | 2.406e-05 *** |
| c(condition) | 21146.0 | 21146.0 | 1 | 589.05 | 226.193 | < 2.2e-16 *** |
| c(number of trees):c(condition) | 204.5 | 204.5 | 1 | 589.07 | 2.187 | 0.1397 |
| AIC | 4498.95 | | | | | |
| BIC | 4525.38 | | | | | |
| Log Likelihood | −2243.47 | | | | | |
| Num. obs. | 605 | | | | | |
| Num. groups: ID | 13 | | | | | |
| Var: ID (Intercept) | 32.42 | | | | | |
| Var: Residual | 93.49 | | | | | |

$^{***}p < 0.001$; $^{**}p < 0.01$; $^{*}p < 0.05$

**distance error**

Table 3

| distance error ∼ number of trees + condition + (1\|participant) | | | | | | |
|---|---|---|---|---|---|---|
| | Sum Sq | Mean Sq | Df1 | Df2 | F | p |
| c(number of trees) | 0.3 | 0.3 | 1 | 723.05 | 0.0076 | 0.9306 |
| c(condition) | 13097.1 | 6548.5 | 2 | 723.04 | 178.3301 | <2e-16 *** |
| AIC | 4810.24 | | | | | |
| BIC | 4837.87 | | | | | |
| Log Likelihood | −2399.12 | | | | | |
| Num. obs. | 739 | | | | | |
| Num. groups: ID | 13 | | | | | |
| Var: ID (Intercept) | 21.85 | | | | | |
| Var: Residual | 36.72 | | | | | |

$^{***}p < 0.001$; $^{**}p < 0.01$; $^{*}p < 0.05$

Table 4

| distance error ∼ number of trees ∗ condition + (1\|participant) (w/o body condition) | | | | | | |
|---|---|---|---|---|---|---|
| | Sum Sq | Mean Sq | Df1 | Df2 | F | p |
| c(number of trees) | 0.3 | 0.3 | 1 | 589.05 | 0.0083 | 0.9273 |
| c(condition) | 8609.8 | 8609.8 | 1 | 589.03 | 223.6304 | <2e-16 *** |
| c(number of trees):c(condition) | 39.4 | 39.4 | 1 | 589.04 | 1.0246 | 0.3118 |
| AIC | 3973.91 | | | | | |
| BIC | 4000.34 | | | | | |
| Log Likelihood | −1980.96 | | | | | |
| Num. obs. | 605 | | | | | |
| Num. groups: ID | 13 | | | | | |
| Var: ID (Intercept) | 27.12 | | | | | |
| Var: Residual | 38.50 | | | | | |

$^{***}p < 0.001$; $^{**}p < 0.01$; $^{*}p < 0.05$

**direction error**

Table 5

| direction error ∼ number of trees + condition + (1\|participant) | | | | | | |
|---|---|---|---|---|---|---|
| | Sum Sq | Mean Sq | Df1 | Df2 | F | p |
| c(number of trees) | 572.6 | 572.6 | 1 | 724.10 | 0.1489 | 0.6997 |
| c(condition) | 18003.6 | 9001.8 | 2 | 723.87 | 2.3405 | 0.0970 . |
| AIC | 8193.80 | | | | | |
| BIC | 8221.43 | | | | | |
| Log Likelihood | −4090.90 | | | | | |
| Num. obs. | 739 | | | | | |
| Num. groups: ID | 13 | | | | | |
| Var: ID (Intercept) | 57.16 | | | | | |
| Var: Residual | 3846.03 | | | | | |

$^{***}p < 0.001$; $^{**}p < 0.01$; $^{*}p < 0.05$

Table 6

| direction error ∼ number of trees ∗ condition + (1\|participant) (w/o body condition) | | | | | | |
|---|---|---|---|---|---|---|
| | Sum Sq | Mean Sq | Df1 | Df2 | F | p |
| c(number of trees) | 528.6 | 528.6 | 1 | 590.15 | 0.1405 | 0.7079 |
| c(condition) | 4.4 | 4.4 | 1 | 589.84 | 0.0012 | 0.9727 |
| c(number of trees):c(condition) | 3477.5 | 3477.5 | 1 | 590.05 | 0.9245 | 0.3367 |
| AIC | 6691.18 | | | | | |
| BIC | 6717.61 | | | | | |
| Log Likelihood | −3339.59 | | | | | |
| Num. obs. | 605 | | | | | |
| Num. groups: ID | 13 | | | | | |
| Var: ID (Intercept) | 50.06 | | | | | |
| Var: Residual | 3761.38 | | | | | |

$^{***}p < 0.001$; $^{**}p < 0.01$; $^{*}p < 0.05$

##### Cartesian standard deviation (sd)

Table 7

| Cartesian sd ∼ number of trees + condition + (1\|participant) | | | | | | |
|---|---|---|---|---|---|---|
| | Sum Sq | Mean Sq | Df1 | Df2 | F | p |
| c(number of trees) | 6.87 | 6.87 | 1 | 99 | 1.0133 | 0.3166 |
| c(condition) | 1432.37 | 358.09 | 4 | 99 | 52.8131 | <2e-16 *** |
| AIC | 584.26 | | | | | |
| BIC | 606.36 | | | | | |
| Log Likelihood | −284.13 | | | | | |
| Num. obs. | 117 | | | | | |
| Num. groups: ID | 13 | | | | | |
| Var: ID (Intercept) | 3.94 | | | | | |
| Var: Residual | 6.78 | | | | | |

$^{***}p < 0.001$; $^{**}p < 0.01$; $^{*}p < 0.05$

Table 8

| Cartesian sd ∼ number of trees ∗ condition + (1\|participant) (w/o body condition) | | | | | | |
|---|---|---|---|---|---|---|
| | Sum Sq | Mean Sq | Df1 | Df2 | F | p |
| c(number of trees) | 6.87 | 6.87 | 1 | 84 | 0.9356 | 0.3362 |
| c(condition) | 1323.15 | 441.05 | 3 | 84 | 60.0623 | < 2e-16 *** |
| c(number of trees):c(condition) | 39.48 | 13.16 | 3 | 84 | 1.7923 | 0.1549 |
| AIC | 523.86 | | | | | |
| BIC | 550.30 | | | | | |
| Log Likelihood | −251.93 | | | | | |
| Num. obs. | 104 | | | | | |
| Num. groups: ID | 13 | | | | | |
| Var: ID (Intercept) | 3.74 | | | | | |
| Var: Residual | 7.34 | | | | | |

$^{***}p < 0.001$; $^{**}p < 0.01$; $^{*}p < 0.05$

**distance sd**

Table 9

| distance sd ∼ number of trees + condition + (1\|participant) | | | | | | |
|---|---|---|---|---|---|---|
| | Sum Sq | Mean Sq | Df1 | Df2 | F | p |
| c(number of trees) | 3.149 | 3.149 | 1 | 99 | 1.4516 | 0.2311 |
| c(condition) | 254.392 | 63.598 | 4 | 99 | 29.3198 | 2.921e-16 *** |
| AIC | 450.56 | | | | | |
| BIC | 472.66 | | | | | |
| Log Likelihood | −217.28 | | | | | |
| Num. obs. | 117 | | | | | |
| Num. groups: ID | 13 | | | | | |
| Var: ID (Intercept) | 0.58 | | | | | |
| Var: Residual | 2.17 | | | | | |

$^{***}p < 0.001$; $^{**}p < 0.01$; $^{*}p < 0.05$

Table 10

| distance sd ∼ number of trees ∗ condition + (1\|participant) (w/o body condition) | | | | | | |
|---|---|---|---|---|---|---|
| | Sum Sq | Mean Sq | Df1 | Df2 | F | p |
| c(number of trees) | 3.149 | 3.149 | 1 | 84 | 1.2912 | 0.2591 |
| c(condition) | 229.262 | 76.421 | 3 | 84 | 31.3381 | 1.188e-13 *** |
| c(number of trees):c(condition) | 5.071 | 1.690 | 3 | 84 | 0.6931 | 0.5588 |
| AIC | 410.77 | | | | | |
| BIC | 437.22 | | | | | |
| Log Likelihood | −195.39 | | | | | |
| Num. obs. | 104 | | | | | |
| Num. groups: ID | 13 | | | | | |
| Var: ID (Intercept) | 0.54 | | | | | |
| Var: Residual | 2.44 | | | | | |

$^{***}p < 0.001$; $^{**}p < 0.01$; $^{*}p < 0.05$

**direction circular sd**

Table 11

| direction circular sd ∼ number of trees + condition + (1\|participant) | | | | | | |
|---|---|---|---|---|---|---|
| | Sum Sq | Mean Sq | Df1 | Df2 | F | p |
| c(number of trees) | 1906.6 | 1906.6 | 1 | 99 | 10.7100 | 0.001468 ** |
| c(condition) | 7053.7 | 1763.4 | 4 | 99 | 9.9059 | 8.778e-07 *** |
| AIC | 950.05 | | | | | |
| BIC | 972.15 | | | | | |
| Log Likelihood | −467.03 | | | | | |
| Num. obs. | 117 | | | | | |
| Num. groups: ID | 13 | | | | | |
| Var: ID (Intercept) | 139.11 | | | | | |
| Var: Residual | 178.02 | | | | | |

$^{***}p < 0.001$; $^{**}p < 0.01$; $^{*}p < 0.05$

Table 12

| direction variance ∼ number of trees ∗ condition + (1\|participant) (w/o body condition) | | | | | | |
|---|---|---|---|---|---|---|
| | Sum Sq | Mean Sq | Df1 | Df2 | F | p |
| c(number of trees) | 1906.6 | 1906.56 | 1 | 84 | 10.5117 | 0.001702 ** |
| c(condition) | 6826.8 | 2275.61 | 3 | 84 | 12.5464 | 7.426e-07 *** |
| c(number of trees):c(condition) | 1106.5 | 368.83 | 3 | 84 | 2.0335 | 0.115378 |
| AIC | 835.64 | | | | | |
| BIC | 862.08 | | | | | |
| Log Likelihood | −407.82 | | | | | |
| Num. obs. | 104 | | | | | |
| Num. groups: ID | 13 | | | | | |
| Var: ID (Intercept) | 137.04 | | | | | |
| Var: Residual | 181.38 | | | | | |

$^{***}p < 0.001$; $^{**}p < 0.01$; $^{*}p < 0.05$

### S2 weighting vs performance

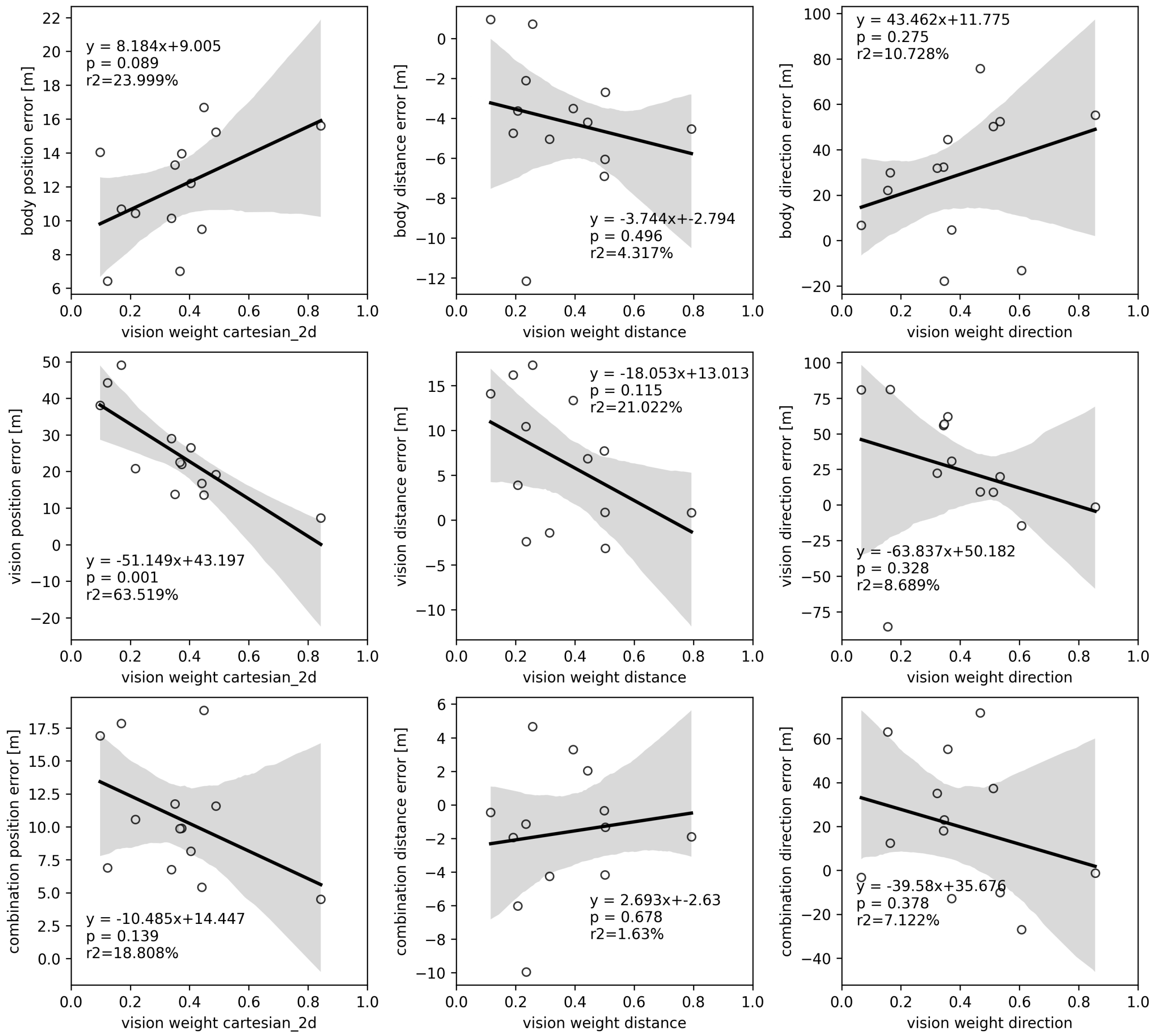


Figure 1: Correlations between observed vision weights in 2D Cartesian (left column), distance (middle column), and direction domain (right column) and position error $[m]$ in body (top row), vision (middle row), and combination condition (bottom row). Regression plots and correlation statistics resemble the content of the main manuscript's Tab. 1. Correlations are described by their $95\% - ci$ (shaded area), regression equation ($y$), p-value ($p$), and coefficient of determination ($r^2$). Data points represent single participants.

### S3 single-cue vs combination biases

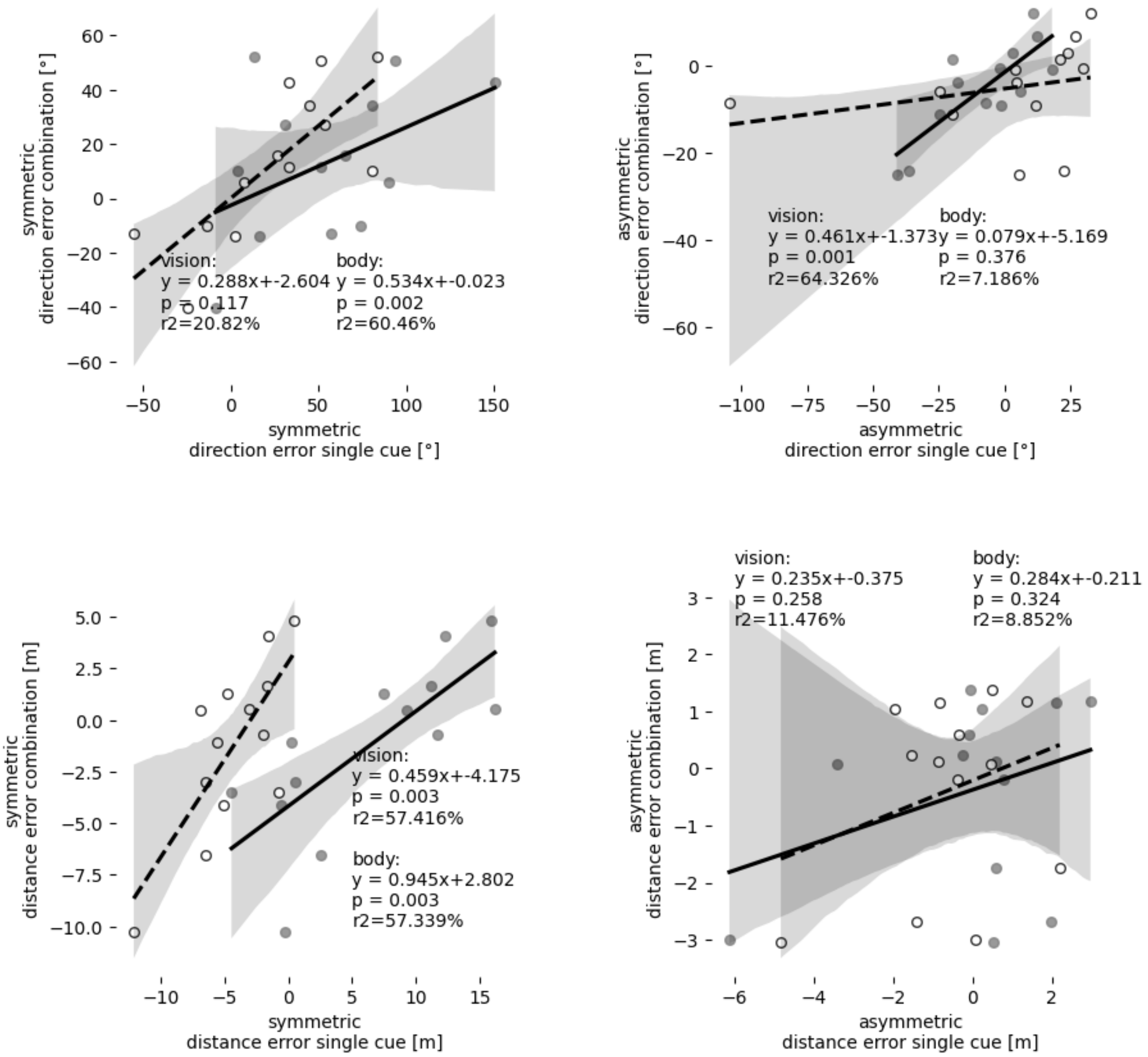


Figure 2: Correlations between symmetric (left column) or asymmetric (right column) error component in direction [°] (top row) or distance domain $[m]$ (bottom row) in either of the single-cue condition (x-axis) and the combination condition (y-axis). Regression plots and correlation statistics resemble the content of the main manuscript's Tab. 2. Correlations are described by their $95\% - ci$ (shaded area), regression equation ($y$), p-value ($p$), and coefficient of determination ($r^2$). Data points represent single participants. Correlations were calculated separately for vision (solid line, filled dots) or body (dashed line, unfilled dots) as the single-cue condition.

### S4 cue weighting vs biases

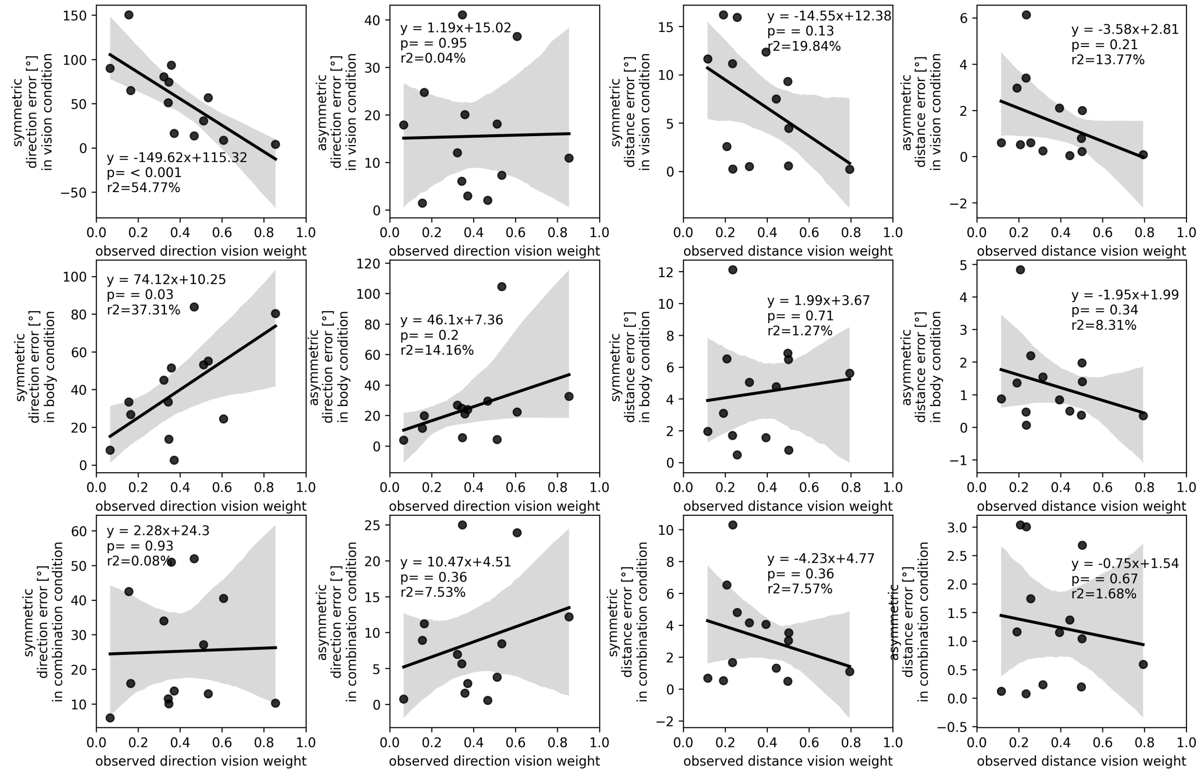


Figure 3: Only symmetric, not asymmetric, direction error component in vision and body condition correlates with vision weight. Correlations of observed vision weights with symmetric (1st and 3rd column) or asymmetric (2nd or 4th column) error component in the vision (1st row), body (2nd row), or combination condition (3rd row) for the direction (1st and 2nd column) and distance domain (3rd and 4th column). Correlations are described by their $95\% - ci$ (shaded area), regression equation ($y$), p-value ($p$), and coefficient of determination ($r^2$).

### S5 gaming experience

Table 13: Categories of gaming experience in hours per week during the most active time on a 5-level scale (levels 1-5 $\sim$ 0, 1-2, 2-4, 4-6, >6 *hours/week*) are correlate with average position error or vision weight across participants. Position error is the mean of body, vision, and combination condition in 0 and 400 tree environments. Observed vision weight is the mean of the 2D Cartesian, distance, and direction domain.

| | | | correlation with: | |
|---|---|---|---|---|
| gaming experience category | mean | sd | position error | observed vision weight |
| (0, 1-2, 2-4, 4-6, >6 *h/week*) | | | $p$ | $p$ |
| 2D | 2.7 | 1.7 | 0.072 | 0.108 |
| 3D | 2.3 | 1.8 | 0.148 | 0.136 |
| 1st person | 2.0 | 1.5 | 0.136 | 0.051 |
| 3rd person | 1.8 | 1.4 | 0.776 | 0.682 |
| with minimap | 2.1 | 1.7 | 0.194 | 0.072 |
| PC | 2.7 | 1.7 | 0.662 | 0.421 |
| console | 2.1 | 1.4 | 0.324 | 0.88 |
| handheld | 2.0 | 1.5 | 0.055 | 0.153 |
| VR | 1.2 | 0.6 | 0.782 | 0.982 |
| average | 2.1 | 1.2 | 0.147 | 0.184 |
